# How to build dense cities without jeopardizing biodiversity? Insights from bird observations in Gothenburg, Sweden

**DOI:** 10.64898/2026.03.03.709206

**Authors:** Ahmed Hazem Eldesoky, Oskar Kindvall, Meta Berghauser Pont, Ioanna Stavroulaki, Jorge Gil, Nadia Charalambous

## Abstract

A central challenge for sustainable urban development is balancing the trade-offs of high urban densities. We examined how biodiversity can be supported within dense or compact urban areas by analyzing bird species richness and composition across 30 sites representing three dense and compact urban form types and species-rich reference areas in Gothenburg, Sweden. Species richness and composition differed significantly between dense-compact urban forms and reference areas, but not between the urban form types themselves. Within dense-compact urban forms, however, sites with higher local natural habitat area and/or better connectivity to surrounding natural habitats supported higher species richness, whereas variation in species composition was associated primarily with local habitat area. Rarely recorded species in dense or compact urban areas were also observed where suitable habitat conditions existed. These results suggest that biodiversity can be supported within dense or compact cities through both site-level interventions and broader-scale planning for habitat connectivity.

## 1. Introduction

To achieve global sustainability goals, urban planners and decision-makers worldwide have adopted strategies that promote denser and more compact urban areas. While higher densities can have positive effects, for example, on public infrastructure, transport, and the economy, they also have several negative environmental, social, and health impacts^1^. Among these negative impacts is the loss of biodiversity within cities, which is crucial for human health and well-being^2^, and has now become a legally binding and urgent priority in some regions, with the enforcement of new policies and legal frameworks, such as the EU Nature Restoration Law. The latter mandates EU cities to restore biodiversity and degraded ecosystems and sets specific targets for urban ecosystems, including no net loss of green urban space and tree cover by 2030, and a steady increase in their total area thereafter^3^. Therefore, a central challenge for sustainable urban development is to identify ways to build dense urban areas without compromising biodiversity.

In urban design and planning, understanding how dense or compact urban development influences biodiversity and how its negative effects can be mitigated remains limited for two main reasons. First, the large majority of studies on urban biodiversity focus on studying variations along the urban-rural gradient, with a lack of studies focusing on intra-urban variations in biotic conditions, especially in dense urban areas^4,5^. Second, although a number of studies (e.g.,^6,7^) have investigated the relationship between urban density and biodiversity within cities, most rely on one-dimensional measures of density such as population density (e.g., people/km^2^), built intensity (e.g., Floor Space Index, FSI) or ground coverage (e.g., Ground Space Index, GSI). One-dimensional measures of density offer limited insights for urban design and planning practice because they are poor indicators of urban form^1,8^. For example, at the same population density, FSI, or GSI, one can still design urban environments with very different morphological characteristics, also creating different social, economic, and environmental conditions^10–12^.

Berghauser Pont and Haupt^10^ have proposed the combination of multiple density measures (e.g., GSI and FSI) in a multi-dimensional categorical classification to measure urban density in ways that better describe urban form and performance. This approach has been shown to be effective in numerically distinguishing between urban form types with distinct morphological characteristics, such as high-rise buildings with little ground coverage and mid-rise perimeter building blocks with relatively high ground coverage, even when they have the same FSI^14^.

In this study, we use this approach to advance understanding of how urban density influences biodiversity by examining the impact of different dense and compact urban form types—defined by different combinations of GSI and FSI—on avian communities in Gothenburg, Sweden. The specific focus on avian communities is because birds are considered good surrogates of overall biodiversity in regions where they are speciose^15^. Furthermore, birds provide key ecosystem services (e.g., pollination, seed dispersal, pest control, and cultural ecosystem services) and are sensitive to changes in the ecosystem, making them excellent study subjects in many areas of ecology^16^. Birds are also widely distributed, highly diverse, and relatively easy to detect and identify^17^.

More specifically, we formulated three hypotheses to investigate how dense or compact urban forms influence two complementary aspects of avian diversity^18^: alpha diversity (i.e., the number of species or species richness within a site) and beta diversity (i.e., variation in species composition among sites).

Our first hypothesis (H1) was that, consistent with prior studies, dense and compact urban forms would exhibit lower species richness and different species composition compared to sparsely built urban forms or semi-natural urban areas (hereinafter referred to as reference areas).

The second hypothesis (H2) was that, within the same dense or compact urban form type, there would be some variation in species richness and composition among sites, but that different dense or compact urban form types would differ significantly in their species richness and composition.

Finally, our third hypothesis (H3) was that variations in species richness and composition in the dense and compact urban forms are associated with site-specific and/or larger-scale environmental characteristics that urban designers and planners can act upon when planning and designing more biodiversity-friendly cities. Examples of key site-specific and larger-scale environmental variables, which are well-documented to influence bird species richness and composition, include local habitat area and diversity, as well as landscape connectivity to surrounding suitable habitat types^4,19–21^.

To test the aforementioned hypotheses, we analyzed bird species occurrence data collected during spring 2024 across 30 urban sites representing three dense and compact urban form types, as well as reference areas in Gothenburg, Sweden^22^. The dense and compact urban form types were identified by Berghauser Pont et al.^23^ based on a statistical clustering of GSI and FSI values across five European cities, including Gothenburg, which also resulted in other urban form types. Here, we use the three most dense and compact types—typical for many European urban cores and consistent with UN-Habitat’s advocacy for compact, high-density development^24^—namely: (1) compact low-rise, (2) compact mid-rise, and (3) dense mid-rise buildings. These three types are distinct in terms of ground coverage (i.e., GSI) or the relationship between built and non-built space (spacious, dense, compact) and building height (low, mid, high). The compact low-rise and compact mid-rise types have similar GSI values but differ in FSI due to differences in building height. The dense mid-rise has the highest GSI and FSI of the three types and is dominated by late 19th-century perimeter building blocks. A detailed description of these types can be found in Berghauser Pont et al.^23^.

## 2. Results

Across the 30 urban sites monitored during spring 2024 (from April 21 to June 16), a total of 61 bird species, representing 25 families, were observed^22^. Table S1 lists all observed bird species, and Figure 1 shows the spatial distribution of the 30 monitoring sites representing the three urban form types (19 sites) and the reference areas (11 sites). More details about how these sites were selected and sampled are provided in Section 4.2. In the following sections, we present the results of the different analyses conducted to test the three hypotheses.

**Figure 1.**
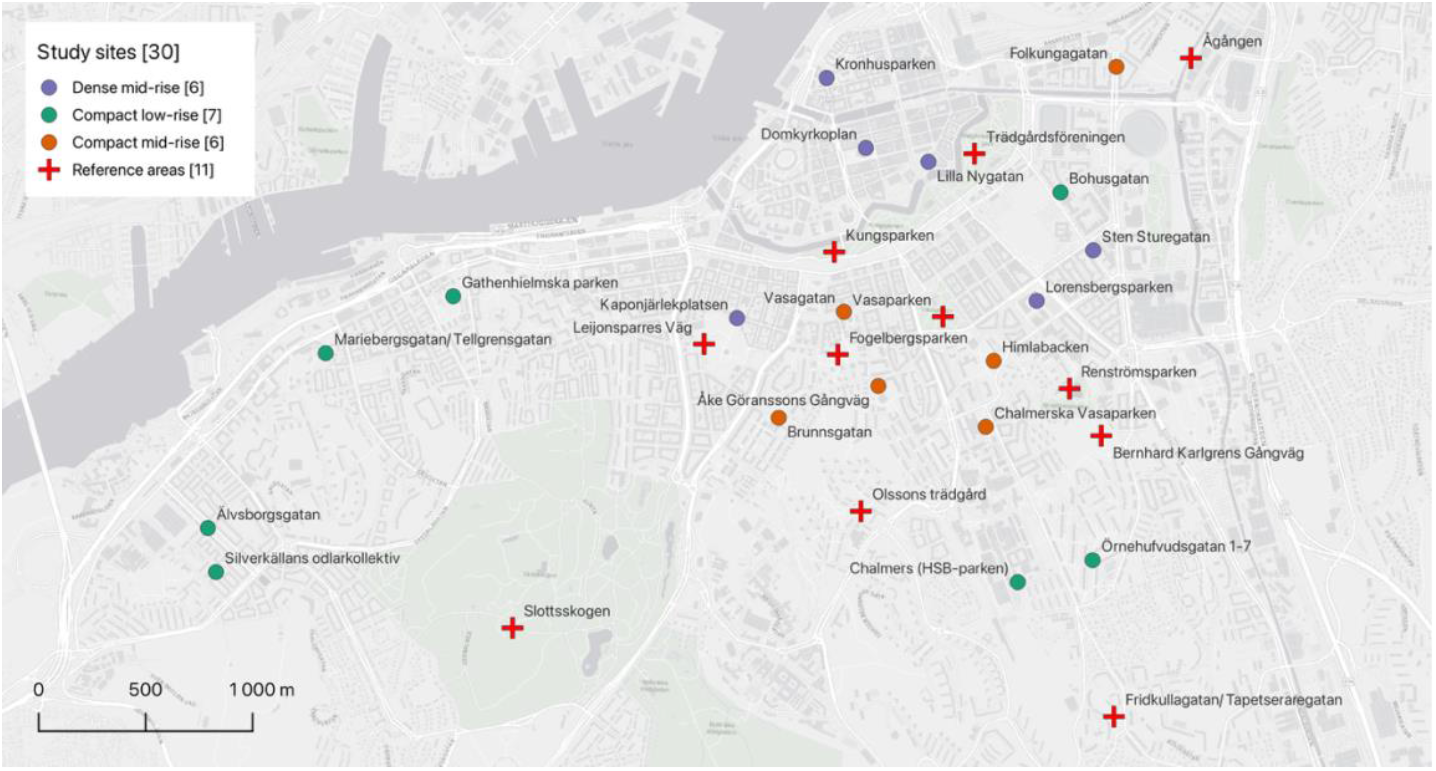
The spatial distribution of the 30 monitoring sites across the study area in Gothenburg, Sweden. Readapted from Eldesoky et al.^22^.

### 2.1. Differences in bird species richness and composition between dense–compact urban forms and reference areas (H1)

To test whether species richness and composition differed between dense–compact urban forms and reference areas (H1), we compared the two groups using the Mann–Whitney U test and permutational multivariate analysis of variance (PERMANOVA), respectively.

Figure 2 shows that species richness was significantly lower in the dense–compact urban forms than in the reference areas (median = 24 vs. 33 species; W_Mann–Whitney_ = 27, *p* < 0.001).

**Figure 2.**
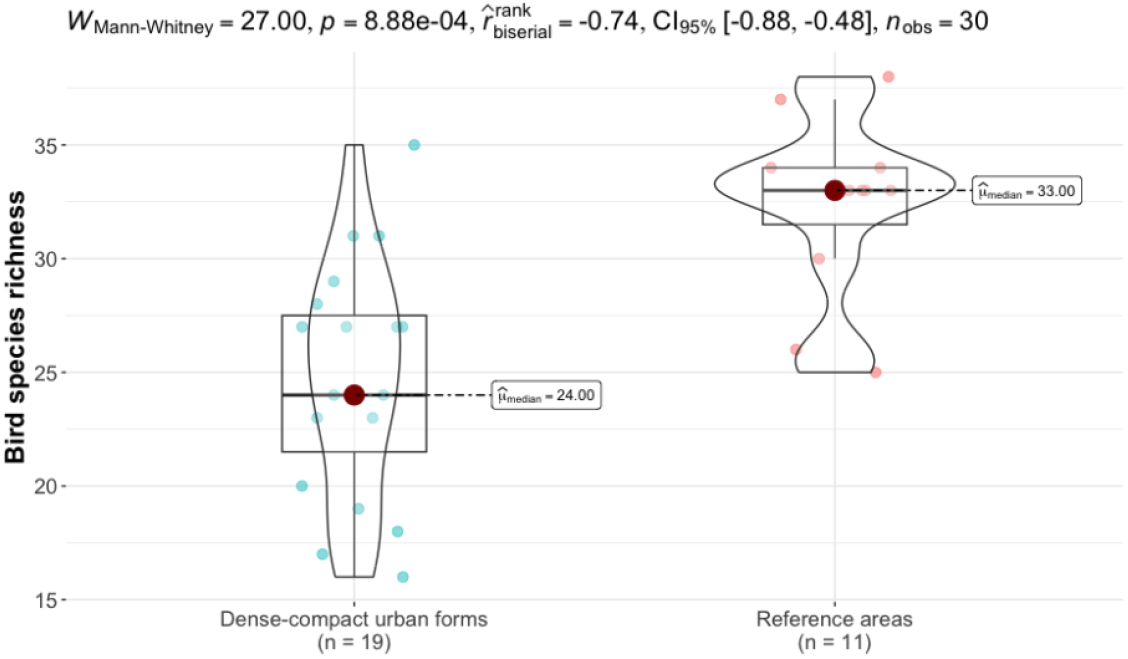
Box and violin plots showing differences in bird species richness between dense–compact urban forms (left) and reference areas (right). Points represent observed values at each site; solid red points denote group medians, and boxes show the interquartile range (IQR).

Similarly, PERMANOVA results showed that there were statistically significant differences in bird species composition between the dense–compact urban forms and reference areas (F = 4.16, R^2^ = 0.13, *p* < 0.001). However, permutational multivariate analysis of dispersion (PERMDISP) showed that the two groups also differed significantly in their within-group multivariate dispersion (*p* = 0.007). This means—given the unbalanced design (19 dense–compact vs. 11 reference sites)—that the significant PERMANOVA result may be driven by differences in centroid location, differences in dispersion, or a combination of both. The non-metric multidimensional scaling (NMDS) ordination (Figure 3), which visualizes relative dissimilarities among sites as distances in a 2D space, supports both effects: the two groups show a shift in centroid position (a location effect), while the dense–compact group shows a more dispersed cloud of points (a dispersion effect). PERMANOVA results from 1000 bootstrap resamples—used to obtain test statistics based on balanced designs, for which PERMANOVA is largely unaffected by heterogeneity—remained consistently significant (*p*-values had a median of 0.002, an IQR of 0.003, and ranged from < 0.001 to 0.041). This further suggests that dense–compact urban forms and reference areas differ in their overall bird species composition. Figure S1 shows a species-per-site presence/absence matrix, where, indeed, several species observed were expected to occur only in either reference areas or in dense–compact settings. For example, the lesser black-backed gull (*Larus fuscus*) was only observed in dense-compact urban forms and not in any of the reference areas, which is expected as the species commonly uses building rooftops for nesting. Similarly, several bird species observed only in reference areas, such as the common reed warbler (*Acrocephalus scirpaceus)* and the European nightjar (*Caprimulgus europaeus*), are not expected to be found in urban environments.

**Figure 3.**
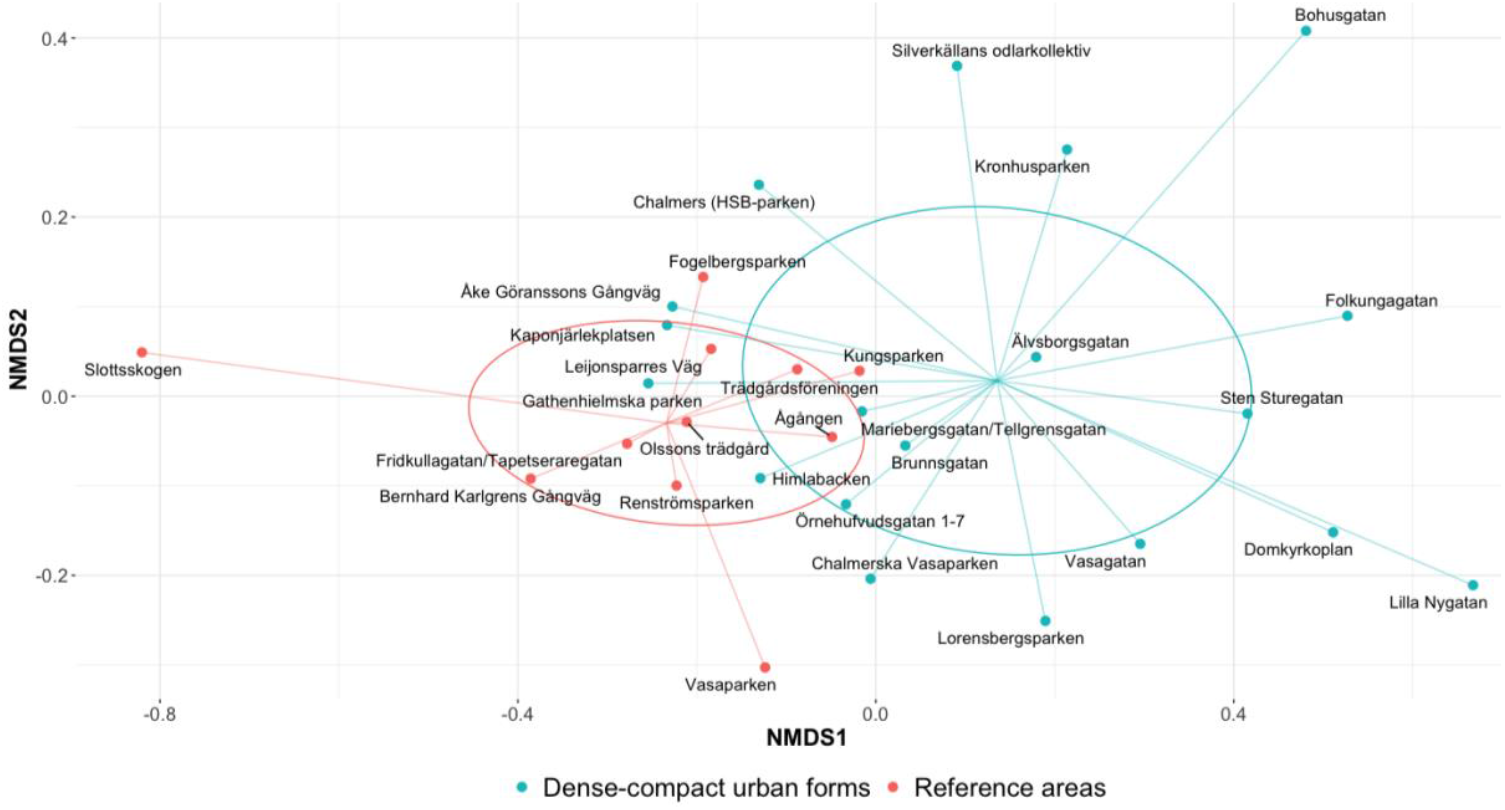
2D-NMDS ordination based on Jaccard dissimilarity. Points represent individual sites belonging to either the dense–compact urban forms (cyan) or the reference areas (red); ellipses show the standard-deviation dispersion of each group; and spider lines connect sites to their group centroid.

Overall, these results, together with the significant differences found in species richness, support H1 and align with previous studies showing that higher urban density is associated with changes in bird species richness and composition.

### 2.2. Bird species richness and composition within and between the dense and compact urban form types (H2)

We next examined variation in species richness and composition within each of the three dense and compact urban form types and tested for significant differences between them (H2). As shown in Figure 4, species richness varied within each urban form type, as indicated by differences in the interquartile range (IQR) and overall range. Among the three types, the compact mid-rise (middle plot) exhibited the widest within-group variation (IQR = 6, range = 13), followed by the compact low-rise (IQR = 4.5, range = 19) and dense mid-rise (IQR = 4, range = 14). However, a Kruskal–Wallis test revealed no significant differences in species richness between the three urban form types (χ^2^ = 1.12, df = 2, *p* = 0.57).

**Figure 4.**
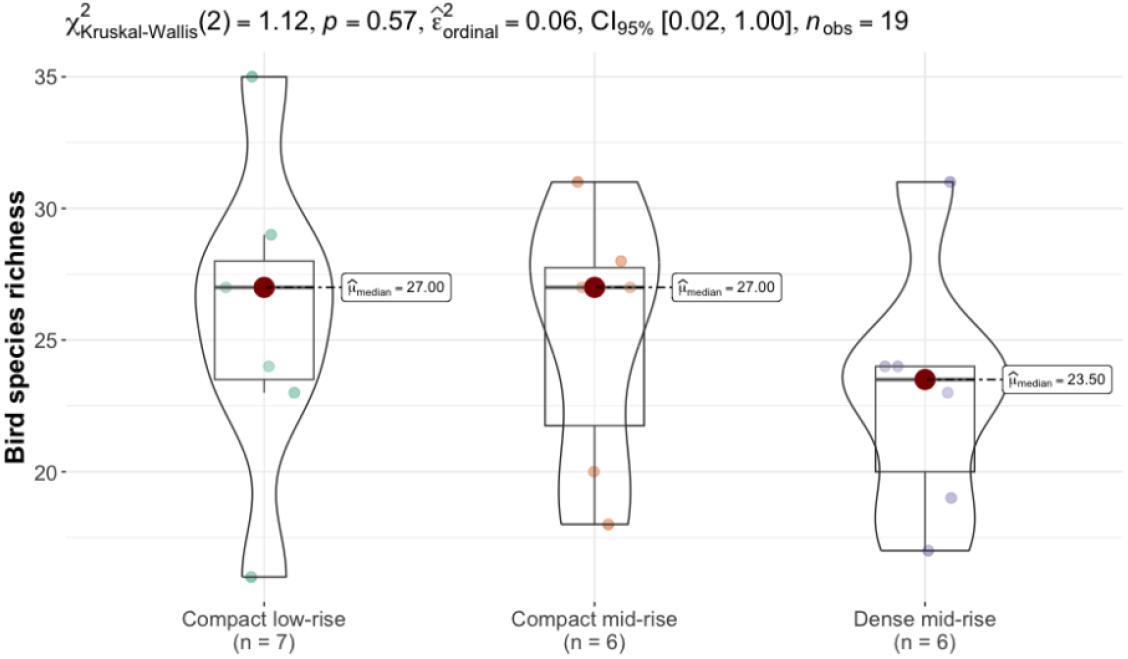
Box and violin plots showing bird species richness across the three dense and compact urban form types: compact low-rise (left), compact mid-rise (middle), dense mid-rise (right). Points represent observed values at each site; solid red points denote group medians; and boxes show the IQR.

As for species composition, the NMDS ordination (Figure 5) shows that sites within each urban form type occupy different locations on the plot. In NMDS, these relative positions reflect differences in species composition. However, the three urban form types also exhibited strong overlap in ordination space, with no clear separation between them.

**Figure 5.**
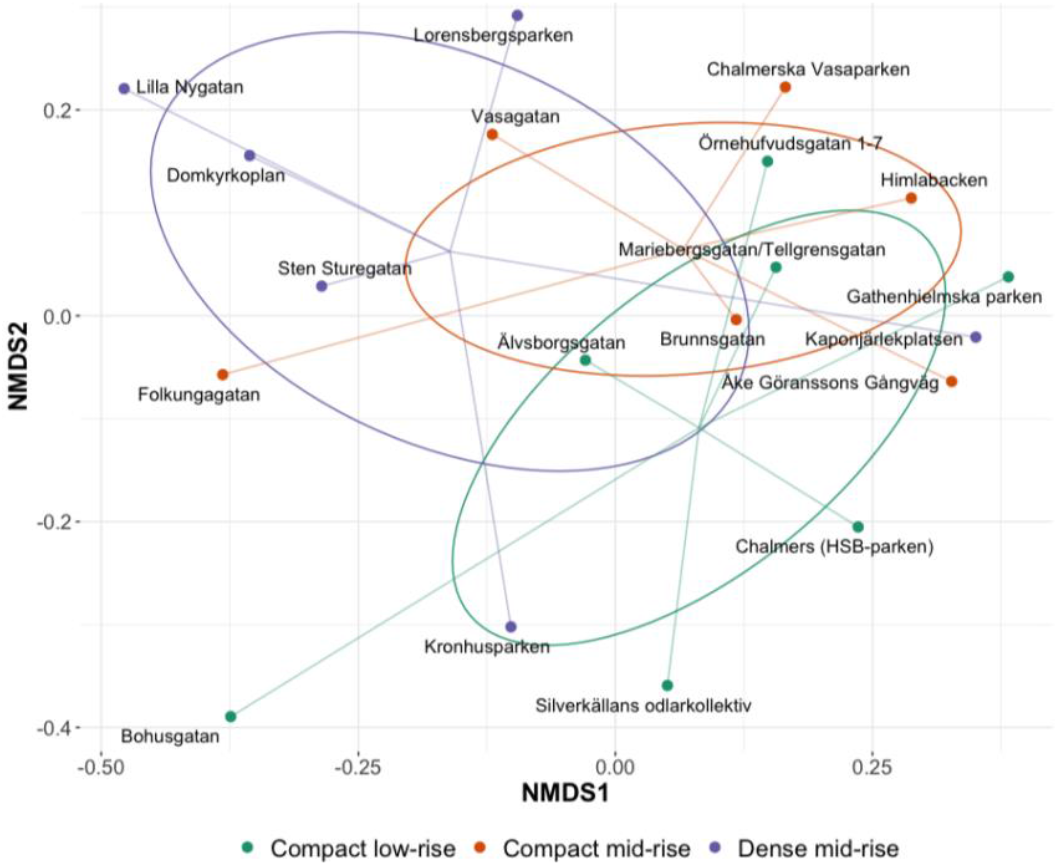
2D-NMDS ordination based on Jaccard dissimilarity. Points represent individual sites belonging to the three dense and compact urban form types (compact low-rise, green; compact mid-rise, orange; dense mid-rise, purple); ellipses show the standard-deviation dispersion of each group; and spider lines connect sites to their group centroid.

The PERMANOVA test showed an overall borderline significant difference in species composition between the three urban form types (F = 1.77, R^2^ = 0.18, *p* = 0.049), and the PERMDISP test showed that the types did not differ significantly in their within-group multivariate dispersion (*p* = 0.94). However, none of the pairwise comparisons between types were found to be significant after correction for multiple testing (BH-adjusted *p*-values: 0.09–0.52). The species-per-site presence/absence matrix (Figure S1) shows that several species seemingly associated with particular urban form types were observed only at single sites within those types. This pattern further suggests that the borderline significant differences detected in the PERMANOVA are likely driven by site-specific characteristics or chance observations rather than by the urban form types themselves. For example, the common tern (*Sterna hirundo*), a water-dependent species, was observed only at a dense mid-rise site (Kronhusparken) located near the Göta älv river (see Figure 1). Similarly, the stock dove (*Columba oenas*), which is rarely recorded in urban environments, was observed only in a green space (Chalmerska Vasaparken) surrounded by compact mid-rise buildings. This particular site has a couple of valuable old beech trees (*Fagus sylvatica*) that provide highly suitable nesting cavities for the species, but also foraging opportunities.

Overall, these findings suggest that, similar to species richness, species composition varied within each of the urban form types but did not differ significantly between them; hence, H2 was not supported.

### 2.3. Relationship between environmental variables and bird species richness and composition within dense–compact urban forms (H3)

To assess whether variation in bird species richness and composition in dense–compact urban forms was associated with site-specific and larger-scale environmental variables (H3), we fitted two sets of models. Specifically, we ran 75 Conway-Maxwell Poisson generalized linear models (GLMs) for species richness and 75 PERMANOVA models for species composition, using all possible combinations of three environmental variables. Briefly, local-scale variables included the area and diversity (based on the Shannon diversity index) of suitable natural habitat types within a 70 m buffer around each site, whereas the larger-scale variable included structural connectivity, measured as the average Euclidean distance to surrounding suitable natural habitats, within multiple buffer distances (200–4000 m) to assess sensitivity to spatial scales. These variables and their calculations are described in detail in Section 4.3.

Among the fitted GLM models, the model combining local habitat area and structural connectivity measured at 1800 m yielded the lowest AICc (Akaike information criterion corrected for small sample size), indicating the most parsimonious model. The model combining local habitat area and connectivity at 1600 m also had strong support (ΔAICc = 1.26). These two models exhibited better relative fit compared to all other candidate models (pseudo-R^2^ = 0.81–0.82), and diagnostic checks (Figure S2) confirmed that the model assumptions were met and supported the adequacy of the most parsimonious model. More specifically, residuals showed no systematic patterns and closely followed the expected theoretical distribution, and there was no evidence of residual spatial autocorrelation (Moran’s I = 0.04, *p* = 0.15).

In both models, local habitat area and structural connectivity revealed statistically significant positive associations with species richness (*p* < 0.001), with habitat area showing a stronger association (Table 2; Figure 6). Specifically, based on the most parsimonious model, a one–standard-deviation increase in local habitat area was associated with 16% (12–21%) increase in expected species richness (IRR = 1.16, 95% CI [1.12, 1.21]) after accounting for structural connectivity, whereas a one–standard-deviation increase in structural connectivity was associated with 10% (5–15%) increase (IRR = 1.10, 95% CI [1.05, 1.15]).

**Table 2.** Best-performing candidate models (ΔAICc < 2). IRR = Incidence Rate Ratio (exponentiated estimate); CI = confidence interval. A detailed description of all fitted models, including standardized estimates, standard errors, z-statistics, pseudo-R^2^, and AICc values, is provided in Table S2.

| Model ID | Variables | IRR | 95% CI for IRR | $p$-value | Pseudo $R^2$ | AICc |
| --- | --- | --- | --- | --- | --- | --- |
| 27 | Intercept | 24.3 | [23.3, 25.4] | $< 2e-16$ | 0.81 | 95.13 |
| | Habitat area | 1.16 | [1.11, 1.21] | $3.28e-12$ | | |
| | Connectivity (1600 m) | 1.09 | [1.05, 1.14] | $6.60e-05$ | | |
| 28 | Intercept | 24.3 | [23.4, 25.3] | $< 2e-16$ | 0.82 | 93.87 |
| | Habitat area | 1.16 | [1.12, 1.21] | $5.36e-13$ | | |
| | Connectivity (1800 m) | 1.10 | [1.05, 1.15] | $2.05e-05$ | | |

**Figure 6.**
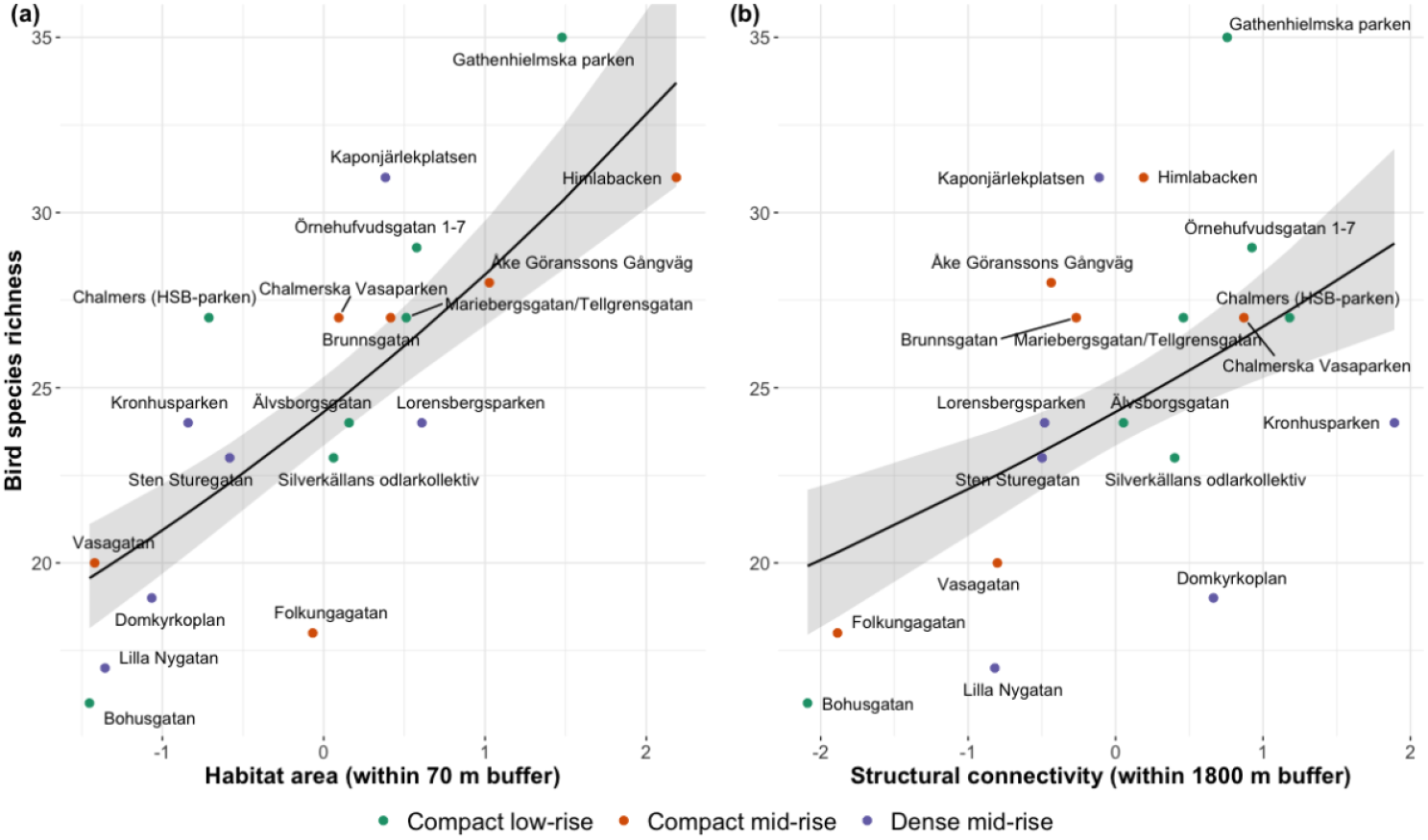
Marginal relationships between bird species richness and (a) local habitat area (within 70 m buffer) and (b) structural connectivity (within 1800 m buffer). Points represent observed values; the solid line shows the model-predicted mean from the fitted most parsimonious model, and the shaded ribbon indicates the 95% confidence interval. Habitat area and structural connectivity were standardized using z scores before modeling.

Across all fitted models, only structural connectivity measured between 1400–2400 m showed statistically significant relationships with species richness when combined with habitat area (*p*-values ranged from < 0.001 to 0.024), with connectivity at 1600 m and 1800 m yielding the strongest associations and lowest AICc values (Table S2). Unlike previous studies (e.g.,^6,19^), the Shannon diversity index did not show a statistically significant relationship with species richness in any models that included local habitat area (Table S2). Although it was significant in models incorporating connectivity measured at 1600, 1800, or 2000 m, those models had ΔAICc > 10, indicating essentially no support. This is likely because, in our case, the Shannon index exhibited little variation across the three urban form types (Figure S3) and thus provided minimal improvement in model fit.

As for species composition, PERMANOVA results (Table S3), assessing the marginal relationship between each of the three variables and species dissimilarity, showed that local habitat area was the only variable significantly associated with species dissimilarity in all models (marginal R^2^ = 0.12–0.22, *p-*values ranged from < 0.001 to 0.015). In contrast, connectivity was not significant in any of the models that included habitat area (*p* > 0.05), indicating that habitat area is the primary factor associated with variation in species composition. Similar to the results for species richness, the Shannon diversity index showed no significant marginal association with species composition (*p* > 0.05).

Overall, these results support H3 that variation in bird species richness and composition in dense and compact urban forms is associated with site-specific and/or larger-scale environmental variables.

## 3. Discussion

A central challenge in sustainable urban development is balancing the benefits and costs of high urban densities. In this study, we examined whether there is a potential to support biodiversity within dense or compact cities. We argued that understanding how high urban density influences biodiversity has been limited in the scientific agenda for several reasons, most importantly because research has largely focused on urban–rural gradients and because urban density is often measured in one-dimensional ways that do not reflect aspects of urban form and performance.

To take a step towards a more nuanced understanding of the relationship between dense and compact urban development and biodiversity, we tested different hypotheses using data on bird species distributions across 30 sites in Gothenburg, Sweden. These sites represented three distinct dense and compact urban form types—defined by different combinations of GSI and FSI—as well as sparsely built urban forms and semi-natural urban areas (referred to as reference areas).

Consistent with prior studies, our findings show that species richness and composition differed significantly between dense–compact urban forms and reference areas (H1). However, there was no significant difference in species richness or composition between the different dense and compact urban form types (H2). Andersson and Colding^25^ found similar results when they investigated the influence of three suburban urban form types on breeding bird diversity in Stockholm, Sweden.

Most importantly, we found that within dense and compact urban forms, sites with higher local natural habitat area and/or better connectivity to surrounding natural habitats (within 1.6–1.8 km) supported higher bird species richness, whereas variation in species composition among sites was driven primarily by local habitat area (H3). Notably, bird species rarely recorded in urban environments (e.g., *Columba oenas*) were observed at certain sites within the dense and compact urban forms where favorable conditions for their existence were present (e.g., suitable foraging and nesting opportunities).

These findings have several important implications for urban design and planning. First, they highlight that expanding the area of natural habitats remains the most effective strategy for supporting species richness and shaping species composition within dense and compact urban areas. As McDonald et al.^8^ demonstrated, creating dense or compact urban areas that share space with nature is possible through a set of interventions that account for urban form as well as the political (e.g., zoning and building rules), social (e.g., community engagement), and economic (e.g., availability of funding) context.

Second, when such interventions are constrained, for example by limited space and competing land-use demands, there remains an opportunity to support more species through broader-scale planning interventions that improve habitat connectivity. In dense or compact cities, such connectivity can be realized, for example, by repurposing existing linear elements in the city (e.g., riverbanks and railway embankments) as well as smaller public and semi-private spaces (e.g., parks, yards, green roofs, vacant land), which can collectively function as ecological stepping stones across the city^26^.

Third, the results suggest that ecologically important species that contribute to maintaining biodiversity or species of conservation interest can still be supported in dense and compact urban areas through targeted interventions that create favorable conditions for them to thrive, such as suitable microhabitats, foraging opportunities, or nesting sites. Such species-focused interventions can serve different objectives, for example: (1) promoting local (site-level) biodiversity by creating ecological conditions that support multiple species; (2) supporting targeted species or species groups of conservation interest by creating the specific ecological conditions they require; and (3) enhancing overall biodiversity across the dense or compact urban area by achieving complementarity among sites, whereby different sites support species not accommodated elsewhere. In urban design and planning practice, achieving these objectives can be operationalized, for example, by placing reproduction or foraging habitats at strategic locations (e.g., areas that facilitate species movement), as suggested by Kindvall et al.^27^. When doing so, it is important to prioritize habitats that support native species rather than non-native ones that may have negative impacts on biodiversity^28^ or pose health risks to humans^29^.

These findings also contribute to the broader “land sharing vs. land sparing” debate in urban ecology^30^. Several previous studies have suggested that land sparing—characterized by dense or compact development that enables the preservation of larger contiguous areas of natural habitat—reduces biodiversity within the dense or compact urban area but helps maintain higher overall biodiversity at urban fringes^31–33^. Our results support this general pattern but provide a more nuanced understanding by showing that, under a land sparing approach, biodiversity in the dense or compact urban area can still be promoted through a careful multi-scale approach that considers both site-specific and broader landscape characteristics. This is important not only for biodiversity conservation but also for sustaining key ecosystem services that benefit urban residents (e.g., health and well-being).

However, it should be noted that the extent to which biodiversity can be promoted within dense or compact urban areas will depend on the species (or higher taxon), as different taxa respond differently to varying levels of urbanization^30,34^. The latter highlights that understanding how different taxa respond to urbanization is an essential aspect to support biodiversity in cities. This is because biodiversity is a multifaceted concept that encompasses several aspects, including, for example, the number of species (richness), the identity of species present, patterns of dominance and rarity, and the functional roles species play within ecosystems^35^.

In this study, although we focused on a couple of important aspects of biodiversity, namely species richness and composition, we examined only one taxonomic group, birds. However, as mentioned earlier, birds often perform well as biodiversity surrogates, particularly in environments where they are relatively speciose^15^. Nonetheless, a more nuanced understanding of the relationship between high urban density and biodiversity would be strengthened by collecting data from additional taxa and by examining other aspects of biodiversity and the urban environment. Here, we considered only three design variables of the urban environment that urban designers and planners can act upon as a starting point, but the list is far from exhaustive. Other factors, such as microclimate, noise, light pollution, and green space management practices, may interact with higher urban density in complex ways that require a more comprehensive understanding of how multiple design, abiotic, biotic, and management factors jointly influence biodiversity^4^.

Finally, collecting and analyzing additional ecological data from within dense and compact urban areas will be an important next step to provide more robust evidence that can support urban design and planning decisions aimed at promoting biodiversity in dense or compact cities. Such data would help not only to confirm and generalize the significant results found in this study (especially H3), but also to further confirm or reject H2 (i.e., whether the type of dense or compact urban form alone influences biodiversity), for which we did not find significant support.

## 4. Methods

### 4.1. Study area

Gothenburg is located on the west coast of Sweden along the Göta älv river (57°42′ N, 11°58′ E). It is the second-largest city in Sweden, with approximately 608,933 inhabitants in 2024 and a population density of 1,352.7 people per km^2^ (Statistics Sweden; https://www.scb.se/). The city has a maritime temperate climate, characterized by relatively cool summers and mild winters for its latitude. Gothenburg lies within the nemoral vegetation zone, dominated by temperate deciduous forests^36^, where most tree species leaf out in late April or May and begin to defoliate around October.

In recent decades, Gothenburg has experienced growth, with the population expected to reach 713,700 by 2050^37^. In response to this urbanization trend and sustainable development goals defined in Agenda 2030, urban densification has been identified as a central development strategy since 2000^38^. Since 2014, the city has pursued a Green Strategy emphasizing green–blue infrastructure, park accessibility, and ecological connectivity, facing challenges where green–blue corridors intersect dense urban areas. In the comprehensive plan from 2022 and thematic plans developed in parallel, such as the Green Plan, Gothenburg prioritizes promoting densification, climate adaptation, and preserving biodiversity.

As for birds, the subject of this study, Gothenburg hosts a relatively rich avifauna, with more than 350 species recorded in the Swedish Species Observation System from 1746 until now (Artportalen; https://artportalen.se/).

### 4.2. Bird observations

For this study, we used a bird species occurrence dataset comprising 239,597 records of 61 species collected between 21 April and 16 June 2024 across 30 urban sites in Gothenburg^22^. Of the 30 sites, 19 represented the three dense and compact urban form types (6–7 sites per type). The other 11 sites were designated as reference sites to test H1. These reference sites were selected in semi-natural urban areas where bird diversity is expected to be higher, such as large urban parks, woodlands, as well as more spacious, sparsely built urban forms with low GSI and FSI and abundant vegetation.

The dataset was produced based on automated detections of bird vocalizations in 10,691 hours of passive acoustic recordings, focused on periods of peak bird vocal activity (dawn, morning, and evening choruses). In each site, acoustic recorders were attached either to trees in publicly accessible green spaces between buildings or to trees along streets to capture the variety of conditions present within each urban form type. Species identification was performed using a state-of-the-art convolutional neural network (CNN) model, namely BirdNET^39^. The automated detections underwent rigorous validation and post-processing, including a manual review of more than 5,000 records by an expert ornithologist. The dataset is available for download in a standard Darwin Core Archive (DwC-A) format at (https://zenodo.org/records/15490818), and a detailed description of how the data were collected, processed, and validated can be found in Eldesoky et al.^22^.

At each site, bird species richness (alpha diversity) was calculated as the cumulative number of distinct species observed. The dataset was filtered to include only records with an occurrence probability ≥ 0.75. The latter represents the likelihood of a species occurrence, for which Eldesoky et al.^22^ recommended a threshold of 0.75 to ensure the accuracy of the records by balancing sensitivity and specificity. Differences in species composition between every pair of sites (beta diversity) were quantified using the Jaccard similarity^40^, which measures the proportion of shared species relative to the total number of species present across the two sites, based on a species presence–absence matrix.

### 4.3. Environmental variables

Three environmental variables known to influence bird species richness and composition were measured in this study. These included two site-specific and one larger-scale variable: the area and diversity of suitable local habitats (Section 4.3.1) and the landscape connectivity to surrounding suitable habitats (Section 4.3.2).

In this study, we focused specifically on suitable natural habitats because, although a few urban bird species can utilize built structures (e.g., buildings and roads) for foraging or nesting, the large majority rely primarily on green and blue natural habitats (Table S1). Green–blue infrastructure is also the most challenging to maintain or expand in dense and compact urban environments, where space is highly limited, making it essential to understand how these habitats can be managed. Because this study examines overall species richness and composition rather than focal species, all suitable natural habitats were considered of equal value. Different bird taxa have diverse habitat requirements^41^, making it unrealistic to assume that a single habitat type would work for all species present in the urban environment.

#### 4.3.1. Local habitat area and diversity

Local natural habitat area and diversity were quantified using a 70 m buffer around each sampling point. This distance corresponds to the maximum reliable detection range of the acoustic recorders used in the study^22^. Restricting measurements to this buffer ensured that the site-specific variables were quantified at the same scale as the bird detections. Local habitat diversity within each buffer was measured using the Shannon diversity index:

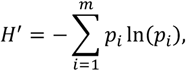

where *p*_*i*_ is the proportion of habitat type *i*.

#### 4.3.2. Landscape connectivity

Landscape connectivity can be measured in different ways, with many different metrics proposed in the literature, which can be very broadly classified into two main groups: structural and functional connectivity^42,43^. Structural connectivity describes the physical configuration of the landscape (i.e., physical arrangement of habitat within the surrounding non-habitat), regardless of the movement capabilities or behavior of the species. It is often measured using Euclidean distance-based metrics^43^. On the other hand, functional connectivity considers species-specific movement abilities and how the landscape either facilitates or impedes movement^44^. In this study, we are primarily interested in studying overall species richness and composition rather than focal species. Therefore, making assumptions about species-level traits (e.g., dispersal ability, behavior, or environmental factors resulting in movement resistance) that could be generalized across the diverse observed bird species may introduce uncertainty and potential errors^44^. Furthermore, dispersal distance is not solely species-specific; it can also vary significantly depending on individual traits such as age and sex^45^. Therefore, for the purpose of this study, we focused on structural rather than functional connectivity.

Specifically, structural connectivity was measured as the inverse of the average Euclidean distance to all suitable natural habitats, calculated within multiple buffer distances (200–4000 m) around each site:

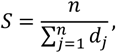

where *n* is the number of suitable habitat pixels, and *d*_*j*_ is the Euclidean distance to the habitat pixel *j*. This measure increases when a site is surrounded by more and/or closer habitats, reflecting higher connectivity, and decreases when habitats are fewer and/or more distant, reflecting lower structural connectivity. Although the measure does not explicitly incorporate habitat area, all habitats in our study were of equal size (10 × 10 m pixels); therefore, the number of habitat pixels represents the amount of surrounding habitat. Consequently, sites with many nearby habitat pixels can be interpreted as being embedded within a larger, more continuous habitat network.

A sensitivity analysis across the different buffer distances was conducted to identify the most appropriate spatial scale (Section 4.4.3).

#### 4.3.3. Biotope map

To calculate the site-specific and larger-scale environmental variables described in the previous sections, we created a biotope map (Figure S4) covering the entire study area and representing all habitat types listed in Table S1. The map was based on the 2023 Swedish National Land Cover Data (NMD2023), freely available from the Swedish Environmental Protection Agency (https://www.naturvardsverket.se/). The base raster layer of NMD2023 has a 10-m spatial resolution and classifies the landscape into 25 habitat classes, of which 16 represent different wet and dry forest types. These forest classes were reclassified into two broader classes more relevant for our analysis of bird communities: Trees (201) and Valuable trees (202). The latter were obtained from a point layer available from the Swedish Species Observation System (Artportalen), which includes trees considered worthy of protection, such as giant, very old, and large hollow trees of native species. We also added a new class for Bushes (200) by combining the class Open land with vegetation (42) in the NMD2023 base raster layer with pixels classified as having vegetation heights between 0.5 and 5 m in the vegetation-height add-on layer included in NMD2023.

Because many bird species nest or forage in habitats adjacent to water, we also defined three beach habitats based on adjacency to water pixels in the NMD2023 base raster layer. These include: Cliff beach (241), Grassland beach (242), and Shaded beach (243). Pixels originally classified as Non-vegetated open land (41) and adjacent to water (61, 62) were reclassified as Cliff beach (241). Similarly, pixels classified as Vegetated open land (42) and adjacent to water were reclassified as Grassland beach (242). Pixels adjacent to water that we initially classified as Bushes (200) or Trees (201) were reclassified as Shaded beach (243). Reeds (444) represent another specific nesting habitat used by some bird species observed in Gothenburg (Table S1). This habitat was identified as pixels in the base NMD2023 raster layer classified as Open wetland (2) and were adjacent to Inland water (61).

In addition to the natural habitat types described above, the biotope map also includes several non-natural, built habitat classes (e.g., buildings, roads, railways, outdoor restaurant/café areas) obtained from official national geographic datasets (Lantmäteriet; https://www.lantmateriet.se/) and OpenStreetMap (OSM). However, as mentioned earlier, although these can be used by some species (Table S1), our analyses focused solely on natural urban habitats, which formed the basis for defining suitable habitats in this study.

### 4.4. Statistical analyses

#### 4.4.1. Testing differences between dense–compact urban forms and reference areas (H1)

To test our first hypothesis that there are statistically significant differences in bird species richness and composition between dense–compact urban forms and reference areas, we used two complementary statistical approaches.

First, differences in species richness between the two groups were assessed using the Mann–Whitney U test^46,47^. This test (also known as the Wilcoxon rank sum test) is a nonparametric alternative to Student’s t-test to assess whether two independent groups differ without requiring the assumption of normality, making it suitable for discrete data (e.g., counts). The analysis was conducted in R (version 4.5.2) using the *rstatix*^48^ and *ggstatsplot*^49^ packages.

Second, differences in species composition were statistically tested using permutational multivariate analysis of variance (PERMANOVA)^50^ and visualized using non-metric multidimensional scaling (NMDS)^51^. PERMANOVA is a non-parametric method for multivariate analysis of variance (ANOVA). It can be used to test whether two or more groups differ significantly in a multivariate response (e.g., species composition), and to partition variation in that response to determine how much variation is associated with different independent variables or predictors. NMDS represents relative dissimilarities among samples as distances in a reduced low-dimensional space (usually 2D). PERMANOVA was performed using the *adonis2* function in the R package *vegan*^52^, and NMDS ordinations were produced using *ggordiplots*^53^. Pairwise dissimilarities between sites were measured using Jaccard dissimilarities (1 – Jaccard similarity^40^), which is appropriate for presence/absence species composition data^54^. All test statistics were computed using 10,000–100,000 permutations, with the number of permutations increased until *p*-values stabilized.

After running PERMANOVA, we tested the homogeneity of multivariate dispersion between groups using permutation tests for homogeneity of multivariate dispersions (PERMDISP). PERMDISP is a multivariate analogue of Levene’s test for homogeneity of variances^55^. In unbalanced designs, such as our comparison of 19 dense–compact and 11 reference sites, differences in dispersion between groups can influence PERMANOVA results^56^. This can lead to PERMANOVA significant outcomes that may reflect differences in dispersion, centroid (location), or a combination of both^54,57,58^ . Therefore, when both PERMANOVA and PERMDISP were significant in the unbalanced design, we examined the NMDS ordination to visually assess whether groups differed mainly in location, dispersion, or both. Furthermore, because PERMANOVA is largely unaffected by heterogeneity for balanced designs^56^, when dispersion was heterogeneous, we reran PERMANOVA using 1000 bootstrap subsamples of the larger group to get balanced designs and obtain more robust test statistics following Sanders-Smith et al.^59^.

#### 4.4.2. Exploring variation within each urban form type and testing differences between them (H2)

For our second hypothesis, we examined whether bird species richness and composition varied within each of the three dense and compact urban form types and differed significantly between them. First, variation in species richness within each urban form type was explored using a combination of box and violin plots. We then used the Kruskal–Wallis test to assess whether species richness differed significantly between the three urban form types. The Kruskal–Wallis test is a nonparametric alternative to one-way ANOVA that assesses whether three or more groups differ based on ranked data^60^. All analyses were performed in R using the *rstatix* and *ggstatsplot* packages.

For species composition, we assessed within-type dissimilarity using NMDS ordination. We then used PERMANOVA to test for statistically significant differences in species composition between the three urban form types. As in H1, PERMANOVA was based on Jaccard dissimilarities and run with 10,000– 100,000 permutations, and homogeneity of multivariate dispersion between urban form types was assessed with PERMDISP prior to interpreting results. Pairwise comparisons between urban form types were conducted using the *pairwise.adonis* function in the *pairwiseAdonis* package^61^, with *p*-values adjusted for multiple comparisons using the Benjamini–Hochberg (BH) correction^62^.

#### 4.4.3. Testing associations with environmental variables (H3)

For our third hypothesis (H3), we tested whether variation in bird species richness and composition in the dense and compact urban forms was associated with site-specific and/or larger-scale environmental variables, introduced in Section 4.3. We evaluated these relationships using two sets of models.

First, to examine whether species richness was associated with local habitat area, the Shannon diversity index, and structural connectivity, we fitted generalized linear models (GLMs). GLMs extend classical linear regression by allowing for non-normal error distributions such as the Poisson distribution, which is appropriate for count data (e.g., species richness). To identify the combination of environmental variables best associated with species richness, we fitted 75 candidate Poisson GLMs representing all possible combinations of the three environmental variables (Table S2). Since structural connectivity was measured at multiple spatial scales (200–4000 m), no two connectivity measures were included in the same model.

Prior to model fitting, we assessed collinearity between variables using pairwise Pearson correlations and excluded variables with correlation coefficients greater than |0.5|^63^. Structural connectivity at 200 m and 400 m had |r| > 0.5 with local habitat area, and therefore excluded (Figure S5).

Poisson GLMs were fitted with log link functions using the base *stats* R package and were checked for potential overdispersion or underdispersion. Many models exhibited significant underdispersion, where the residual variance is smaller than expected under the Poisson assumption (variance = mean). Using a Poisson GLM in such cases can lead to overestimated standard errors and, consequently, misleading inferences^64^. To address this, all candidate models were refitted using the Conway-Maxwell-Poisson distribution, which can handle both underdispersion and overdispersion^65,66^. The Conway-Maxwell-Poisson distribution is available in the *glmmTMB package* (version 1.1.13)^67^ in R, which we used to refit all the GLM candidate models.

The parsimony of the candidate models was evaluated using the Akaike Information Criterion corrected for small sample size (AICc)^68^. We calculated ΔAICc values (relative to the lowest AICc) to identify models with substantial support (ΔAICc < 2). Nagelkerke’s pseudo-R^269^ was used to compare the relative fit of candidate models to the null model. Model assumptions and the adequacy of the most parsimonious model were further assessed using simulation-based residual diagnostics and Moran’s I test for spatial autocorrelation, both implemented in the *DHARMa* package^70^ in R.

Second, to assess how much variation in species composition was associated with each environmental variable, and to test the significance of these associations, we fitted PERMANOVA models with marginal tests. Similar to the species richness analysis, we fitted 75 PERMANOVA models representing all possible combinations of the three environmental variables, using the previously calculated Jaccard dissimilarities in species composition as the response. For each model, we reported the permutation-based *p*-values and the marginal R^2^, which represents the unique contribution of each variable after accounting for the influence of all others.

## 5. Data availability

The bird species occurrence dataset used in this study is freely available on Zenodo (version 2.0.0; https://zenodo.org/records/15490818). The biotope map used to calculate the different environmental variables described in Section 4.3 can be made available upon request as a GeoTIFF raster.

## 6. Code availability

The code used to calculate the environmental variables, conduct all statistical analyses, and produce all figures can be made available upon request.

## 8. Author contributions

M.B.P., A.H.E., I.S., O.K., and J.G. conceptualized the study. A.H.E. analyzed the data, conducted the formal analyses, and prepared the figures. O.K. produced the biotope map and provided ecological insights. A.H.E., M.B.P., O.K., I.S., and J.G. contributed to the interpretation of the results, and N.C. provided insights into their practical applicability. A.H.E. prepared the initial draft with contributions from O.K. and M.B.P. All authors revised and edited the manuscript and approved the final version. M.B.P. and N.C. acquired the funding.

