## Supplementary material for "How to build dense cities without jeopardizing biodiversity? Insights from bird observations in Gothenburg, Sweden": Table S1

**Table S1.** List of all bird species observed in the study, alongside the habitat types included in the biotope map. For each species–habitat type combination, the table indicates whether the habitat is used for nesting (N), foraging (F), or both (NF). Species are sorted according to the taxonomic order provided by the Swedish Species Information Centre (Artportalen).

**Habitat type**

| Scientific name | Family | Common English name | Buildings (51) | Artificial surfaces (52) | Roads and raiways (53) | Cafés and restaurants (500) | Arable land (3) | Non vegetated open land (41) | Grassland and meadows (42) | Bushes (200) | Trees (201) | Valuable trees (202) | Open wetland (2) | Inland water (61) | Marine water (62) | Cliff beach (441) | Grassland beach (442) | Shaded beach (443) | Reed (444) |
|---|---|---|---|---|---|---|---|---|---|---|---|---|---|---|---|---|---|---|---|
| *Branta canadensis* | *Anatidae* | Canada Goose | | | | | F | | F | | | | F | F | F | N | NF | | N |
| *Branta leucopsis* | *Anatidae* | Barnacle Goose | | | | | | | F | | | | | | | N | NF | | |
| *Anser anser* | *Anatidae* | Greylag Goose | | | | | F | | F | | | | F | F | F | N | NF | | N |
| *Anas platyrhynchos* | *Anatidae* | Mallard | | | | | | | F | | | | | F | F | F | NF | N | N |
| *Bucephala clangula* | *Anatidae* | Common Goldeneye | | | | | | | | | | | | F | | | | N | |
| *Caprimulgus europaeus* | *Caprimulgidae* | European Nightjar | | | | | | | | | N | N | | | | | | | |
| *Apus apus* | *Apodidae* | Common Swift | NF | F | F | F | F | F | F | F | F | F | F | F | | F | F | F | F |
| *Cuculus canorus* | *Cuculidae* | Common Cuckoo | | | | | | | F | NF | NF | NF | F | | | | | NF | NF |
| *Streptopelia decaocto* | *Columbidae* | Eurasian Collared Dove | | | | F | | | F | N | N | N | | | | | | N | |
| *Columba livia* | *Columbidae* | Common Pigeon | N | F | F | F | | | F | | | | | | | | | | |
| *Columba oenas* | *Columbidae* | Stock Dove | | | | | | | | | NF | NF | | | | | | NF | |
| *Columba palumbus* | *Columbidae* | Common Wood Pigeon | | | | | F | | F | | N | NF | | | | | | N | |
| *Gallinula chloropus* | *Rallidae* | Common Moorhen | | | | | | | | | | | | F | | | F | N | N |
| *Fulica atra* | *Rallidae* | Eurasian Coot | | | | | | | | | | | | F | F | | F | N | N |
| *Haematopus ostralegus* | *Haematopodidae* | Eurasian Oystercatcher | | | | | | | F | | | | | | F | N | F | | |
| *Sterna hirundo* | *Laridae* | Common Tern | | | | | | | | | | | | F | F | N | | | |
| *Chroicocephalus ridibundus* | *Laridae* | Black-headed Gull | | | | F | | | F | | | | | F | F | NF | F | | N |
| *Larus canus* | *Laridae* | Common Gull | | | | F | F | | F | | | | | F | F | NF | F | | |
| *Larus argentatus* | *Laridae* | European Herring Gull | | | | F | F | | F | | | | | F | F | NF | F | | |
| *Larus fuscus* | *Laridae* | Lesser Black-backed Gull | N | | | F | | | F | | | | | F | F | NF | F | | |
| *Strix aluco* | *Strigidae* | Tawny Owl | N | F | | | | | F | F | F | NF | | | | | | N | |
| *Dendrocopos major* | *Picidae* | Great Spotted Woodpecker | | | | | | | | | NF | NF | | | | | | NF | |
| *Picus viridis* | *Picidae* | European Green Woodpecker | | | | | | | | | NF | NF | | | | | | NF | |
| *Pica pica* | *Corvidae* | Eurasian Magpie | | | | F | | | F | F | N | N | | | | F | F | N | F |
| *Corvus monedula* | *Corvidae* | Western Jackdaw | N | | | F | F | | F | F | F | N | | | | F | F | F | |
| *Corvus cornix* | *Corvidae* | Hooded Crow | | | | F | | | F | F | NF | NF | | | | F | F | NF | |
| *Poecile palustris* | *Paridae* | Marsh Tit | | | | | | | | F | NF | NF | | | | | | N | |
| *Cyanistes caeruleus* | *Paridae* | Eurasian Blue Tit | | | | | | | | F | NF | NF | | | | | | N | |
| *Parus major* | *Paridae* | Great Tit | | | | | | | | F | NF | NF | | | | | | N | |
| *Phylloscopus sibilatrix* | *Phylloscopidae* | Wood Warbler | | | | | | | | N | NF | NF | | | | | | N | |
| *Phylloscopus trochilus* | *Phylloscopidae* | Willow Warbler | | | | | | | | N | NF | NF | | | | | | N | |
| *Phylloscopus collybita* | *Phylloscopidae* | Common Chiffchaff | | | | | | | | N | NF | NF | | | | | | N | |
| *Acrocephalus scirpaceus* | *Acrocephalidae* | Common Reed Warbler | | | | | | | | | | | | | | | | F | NF |
| *Acrocephalus palustris* | *Acrocephalidae* | Marsh Warbler | | | | | | | | NF | | | | | | | | | |
| *Hippolais icterina* | *Acrocephalidae* | Icterine Warbler | | | | | | | | N | NF | NF | | | | | | N | |
| *Sylvia atricapilla* | *Sylviidae* | Eurasian Blackcap | | | | | | | | N | NF | NF | | | | | | N | |
| *Sylvia borin* | *Sylviidae* | Garden Warbler | | | | | | | | N | NF | NF | | | | | | N | |
| *Curruca curruca* | *Sylviidae* | Lesser Whitethroat | | | | | | | | NF | | | | | | | | | |
| *Curruca communis* | *Sylviidae* | Common Whitethroat | | | | | | | | NF | | | | | | | | | |
| *Regulus regulus* | *Regulidae* | Goldcrest | | | | | | | | F | NF | NF | | | | | | N | |
| *Troglodytes troglodytes* | *Troglodytidae* | Eurasian Wren | | | | | | | | N | NF | NF | | | | | | N | |
| *Sitta europaea* | *Sittidae* | Eurasian Nuthatch | | | | | | | | | NF | NF | | | | | | NF | |
| *Certhia familiaris* | *Certhiidae* | Eurasian Treecreeper | | | | | | | | | NF | NF | | | | | | NF | |
| *Sturnus vulgaris* | *Sturnidae* | Common Starling | | | | | | | F | F | NF | NF | | | | F | F | N | F |
| *Turdus philomelos* | *Turdidae* | Song Thrush | | | | | | | F | F | NF | NF | | | | | F | N | |
| *Turdus merula* | *Turdidae* | Common Blackbird | | | | | | | F | N | NF | NF | | | | | F | N | |
| *Turdus pilaris* | *Turdidae* | Fieldfare | | | | | | | F | F | NF | NF | | | | | F | N | |
| *Erithacus rubecula* | *Muscicapidae* | European Robin | | | | | | | | F | NF | NF | | | | | | N | |
| *Ficedula hypoleuca* | *Muscicapidae* | European Pied Flycatcher | | | | | | | | F | NF | NF | | | | | | NF | |
| *Phoenicurus ochruros* | *Muscicapidae* | Black Redstart | N | F | F | | | | | F | F | F | | | | | | | |
| *Phoenicurus phoenicurus* | *Muscicapidae* | Common Redstart | | | | | | | | F | NF | NF | | | | | | N | |
| *Passer montanus* | *Passeridae* | Eurasian Tree Sparrow | N | | | F | F | | | NF | F | F | | | | | | NF | |
| *Passer domesticus* | *Passeridae* | House Sparrow | N | F | F | F | F | | | NF | F | F | | | | | | NF | |
| *Motacilla alba* | *Motacillidae* | White Wagtail | N | F | F | | F | | F | F | F | F | F | | | F | F | | |
| *Fringilla coelebs* | *Fringillidae* | Chaffinch | | | | | | | | NF | NF | NF | | | | | | NF | |
| *Fringilla montifringilla* | *Fringillidae* | Brambling | | | | | | | | NF | NF | NF | | | | | | NF | |
| *Coccothraustes coccothraustes* | *Fringillidae* | Hawfinch | | | | | | | | NF | NF | NF | | | | | | NF | |
| *Chloris chloris* | *Fringillidae* | European Greenfinch | | | | | | | | NF | NF | NF | | | | | | N | |
| *Acanthis flammea* | *Fringillidae* | Redpoll | | | | | | | | NF | NF | NF | | | | | | NF | |
| *Carduelis carduelis* | *Fringillidae* | European Goldfinch | | F | | | F | F | F | NF | NF | NF | | | | | F | NF | |
| *Spinus spinus* | *Fringillidae* | Eurasian Siskin | | | | | | | | NF | NF | NF | | | | | | NF | |
