## Supplementary material for "How to build dense cities without jeopardizing biodiversity? Insights from bird observations in Gothenburg, Sweden": Table S2

Table S2. Results of the Conway–Maxwell–Poisson GLM candidate models examining the relationship between species richness and three environmental variables (habitat area, Shannon index, and structural connectivity at multiple spatial scales). For each variable included in a model, the standardized estimate (± standard error) is reported, followed by the z-value, p-value, and significance level. Variables not included in a model are left blank. The table also reports the model intercept, AICc, Nagelkerke's pseudo-$R^2$, and dispersion estimates (with significance) from a pre-fitted Poisson GLM based on the Cameron & Trivedi (1990) dispersion test. All variables were standardized prior to modeling to allow direct comparison of effect sizes. Significance levels are indicated as: p < 0.05 = *, p < 0.01 = **, p < 0.001 = ***.

| Model ID | Model | Intercept | habitat_area | shannon_index | str_connectivity_600m | str_connectivity_800m | str_connectivity_1000m | str_connectivity_1200m | str_connectivity_1400m | str_connectivity_1600m | str_connectivity_1800m | str_connectivity_2000m | str_connectivity_2200m | str_connectivity_2400m | str_connectivity_2600m | str_connectivity_2800m | str_connectivity_3000m | str_connectivity_3200m | str_connectivity_3400m | str_connectivity_3600m | str_connectivity_3800m | str_connectivity_4000m | Pseudo R2 | Dispersion | AICc | ΔAICc |
|---|---|---|---|---|---|---|---|---|---|---|---|---|---|---|---|---|---|---|---|---|---|---|---|---|---|---|
| 1 | habitat_area | 3.20 (0.03), z = 112.07, p < 0.001 *** | 0.17 (0.03), z = 5.83, p < 0.001 *** | | | | | | | | | | | | | | | | | | | | 0.642 | 0.37, p < 0.001 *** | 103.39 | 9.52 |
| 2 | shannon_index | 3.21 (0.05), z = 69.11, p < 0.001 *** | | -0.02 (0.05), z = -0.51, p = 0.612 | | | | | | | | | | | | | | | | | | | 0.013 | 0.99, p = 0.98 | 122.57 | 28.7 |
| 3 | str_connectivity_600m | 3.21 (0.05), z = 69.38, p < 0.001 *** | | | -0.03 (0.05), z = -0.64, p = 0.520 | | | | | | | | | | | | | | | | | | 0.022 | 1, p = 0.996 | 122.41 | 28.54 |
| 4 | str_connectivity_800m | 3.21 (0.04), z = 71.44, p < 0.001 *** | | | | -0.06 (0.05), z = -1.29, p = 0.197 | | | | | | | | | | | | | | | | | 0.081 | 0.95, p = 0.802 | 121.22 | 27.35 |
| 5 | str_connectivity_1000m | 3.21 (0.05), z = 69.25, p < 0.001 *** | | | | | -0.03 (0.05), z = -0.58, p = 0.559 | | | | | | | | | | | | | | | | 0.018 | 1, p = 0.997 | 122.48 | 28.61 |
| 6 | str_connectivity_1200m | 3.21 (0.05), z = 70.17, p < 0.001 *** | | | | | | 0.04 (0.05), z = 0.95, p = 0.343 | | | | | | | | | | | | | | | 0.045 | 0.98, p = 0.953 | 121.95 | 28.08 |
| 7 | str_connectivity_1400m | 3.20 (0.04), z = 74.95, p < 0.001 *** | | | | | | | 0.09 (0.04), z = 1.98, p = 0.048 * | | | | | | | | | | | | | | 0.171 | 0.86, p = 0.573 | 119.28 | 25.41 |
| 8 | str_connectivity_1600m | 3.20 (0.04), z = 81.53, p < 0.001 *** | | | | | | | | 0.12 (0.04), z = 2.89, p = 0.004 ** | | | | | | | | | | | | | 0.306 | 0.72, p = 0.14 | 115.9 | 22.03 |
| 9 | str_connectivity_1800m | 3.20 (0.04), z = 82.22, p < 0.001 *** | | | | | | | | | 0.12 (0.04), z = 2.95, p = 0.003 ** | | | | | | | | | | | | 0.319 | 0.71, p = 0.112 | 115.55 | 21.68 |
| 10 | str_connectivity_2000m | 3.20 (0.04), z = 76.32, p < 0.001 *** | | | | | | | | | | 0.09 (0.04), z = 2.19, p = 0.029 * | | | | | | | | | | | 0.202 | 0.83, p = 0.342 | 118.55 | 24.68 |
| 11 | str_connectivity_2200m | 3.20 (0.04), z = 75.97, p < 0.001 *** | | | | | | | | | | | 0.09 (0.04), z = 2.13, p = 0.033 * | | | | | | | | | | 0.194 | 0.84, p = 0.353 | 118.74 | 24.87 |
| 12 | str_connectivity_2400m | 3.20 (0.04), z = 76.54, p < 0.001 *** | | | | | | | | | | | | 0.09 (0.04), z = 2.23, p = 0.026 * | | | | | | | | | 0.207 | 0.83, p = 0.349 | 118.44 | 24.57 |
| 13 | str_connectivity_2600m | 3.20 (0.04), z = 76.05, p < 0.001 *** | | | | | | | | | | | | | 0.09 (0.04), z = 2.16, p = 0.031 * | | | | | | | | 0.195 | 0.83, p = 0.351 | 118.7 | 24.83 |
| 14 | str_connectivity_2800m | 3.20 (0.04), z = 75.84, p < 0.001 *** | | | | | | | | | | | | | | 0.09 (0.04), z = 2.13, p = 0.033 * | | | | | | | 0.191 | 0.84, p = 0.35 | 118.81 | 24.94 |
| 15 | str_connectivity_3000m | 3.20 (0.04), z = 75.87, p < 0.001 *** | | | | | | | | | | | | | | | 0.09 (0.04), z = 2.14, p = 0.033 * | | | | | | 0.192 | 0.84, p = 0.359 | 118.79 | 24.92 |
| 16 | str_connectivity_3200m | 3.20 (0.04), z = 76.31, p < 0.001 *** | | | | | | | | | | | | | | | | 0.09 (0.04), z = 2.20, p = 0.028 * | | | | | 0.201 | 0.82, p = 0.325 | 118.56 | 24.69 |
| 17 | str_connectivity_3400m | 3.20 (0.04), z = 76.83, p < 0.001 *** | | | | | | | | | | | | | | | | | 0.09 (0.04), z = 2.28, p = 0.023 * | | | | 0.213 | 0.81, p = 0.278 | 118.29 | 24.42 |
| 18 | str_connectivity_3600m | 3.20 (0.04), z = 77.66, p < 0.001 *** | | | | | | | | | | | | | | | | | | 0.10 (0.04), z = 2.40, p = 0.016 * | | | 0.231 | 0.79, p = 0.231 | 117.85 | 23.98 |
| 19 | str_connectivity_3800m | 3.20 (0.04), z = 78.55, p < 0.001 *** | | | | | | | | | | | | | | | | | | | 0.10 (0.04), z = 2.52, p = 0.012 * | | 0.249 | 0.77, p = 0.194 | 117.4 | 23.53 |
| 20 | str_connectivity_4000m | 3.20 (0.04), z = 78.57, p < 0.001 *** | | | | | | | | | | | | | | | | | | | | 0.10 (0.04), z = 2.52, p = 0.012 * | 0.25 | 0.78, p = 0.21 | 117.38 | 23.51 |
| 21 | habitat_area + shannon_index | 3.19 (0.03), z = 114.87, p < 0.001 *** | 0.17 (0.03), z = 5.99, p < 0.001 *** | 0.03 (0.03), z = 0.99, p = 0.320 | | | | | | | | | | | | | | | | | | | 0.66 | 0.35, p < 0.001 *** | 105.68 | 11.81 |
| 22 | habitat_area + str_connectivity_600m | 3.19 (0.03), z = 119.11, p < 0.001 *** | 0.17 (0.03), z = 6.30, p < 0.001 *** | | -0.04 (0.03), z = -1.59, p = 0.113 | | | | | | | | | | | | | | | | | | 0.684 | 0.33, p < 0.001 *** | 104.26 | 10.39 |
| 23 | habitat_area + str_connectivity_800m | 3.19 (0.03), z = 116.33, p < 0.001 *** | 0.16 (0.03), z = 5.82, p < 0.001 *** | | | -0.04 (0.03), z = -1.23, p = 0.217 | | | | | | | | | | | | | | | | | 0.669 | 0.34, p < 0.001 *** | 105.18 | 11.31 |
| 24 | habitat_area + str_connectivity_1000m | 3.20 (0.03), z = 112.33, p < 0.001 *** | 0.16 (0.03), z = 5.79, p < 0.001 *** | | | | -0.01 (0.03), z = -0.30, p = 0.762 | | | | | | | | | | | | | | | | 0.644 | 0.37, p < 0.001 *** | 106.56 | 12.69 |
| 25 | habitat_area + str_connectivity_1200m | 3.19 (0.03), z = 114.58, p < 0.001 *** | 0.16 (0.03), z = 5.82, p < 0.001 *** | | | | | 0.03 (0.03), z = 0.94, p = 0.348 | | | | | | | | | | | | | | | 0.658 | 0.36, p < 0.001 *** | 105.79 | 11.92 |
| 26 | habitat_area + str_connectivity_1400m | 3.19 (0.03), z = 126.69, p < 0.001 *** | 0.15 (0.03), z = 6.11, p < 0.001 *** | | | | | | 0.06 (0.03), z = 2.34, p = 0.019 * | | | | | | | | | | | | | | 0.722 | 0.3, p < 0.001 *** | 101.88 | 8.01 |
| 27 | habitat_area + str_connectivity_1600m | 3.19 (0.02), z = 150.74, p < 0.001 *** | 0.15 (0.02), z = 6.97, p < 0.001 *** | | | | | | | 0.09 (0.02), z = 3.99, p < 0.001 *** | | | | | | | | | | | | | 0.806 | 0.21, p < 0.001 *** | 95.13 | 1.26 |
| 28 | habitat_area + str_connectivity_1800m | 3.19 (0.02), z = 155.67, p < 0.001 *** | 0.15 (0.02), z = 7.22, p < 0.001 *** | | | | | | | | 0.10 (0.02), z = 4.26, p < 0.001 *** | | | | | | | | | | | | 0.818 | 0.19, p < 0.001 *** | 93.87 | 0 |
| 29 | habitat_area + str_connectivity_2000m | 3.19 (0.02), z = 146.36, p < 0.001 *** | 0.16 (0.02), z = 7.37, p < 0.001 *** | | | | | | | | | 0.09 (0.02), z = 3.72, p < 0.001 *** | | | | | | | | | | | 0.794 | 0.22, p < 0.001 *** | 96.25 | 2.38 |
| 30 | habitat_area + str_connectivity_2200m | 3.19 (0.02), z = 134.87, p < 0.001 *** | 0.16 (0.02), z = 6.62, p < 0.001 *** | | | | | | | | | | 0.07 (0.02), z = 2.96, p = 0.003 ** | | | | | | | | | | 0.756 | 0.26, p < 0.001 *** | 99.43 | 5.56 |
| 31 | habitat_area + str_connectivity_2400m | 3.19 (0.03), z = 125.86, p < 0.001 *** | 0.15 (0.03), z = 5.93, p < 0.001 *** | | | | | | | | | | | 0.06 (0.03), z = 2.26, p = 0.024 * | | | | | | | | | 0.718 | 0.29, p < 0.001 *** | 102.11 | 8.24 |
| 32 | habitat_area + str_connectivity_2600m | 3.19 (0.03), z = 121.25, p < 0.001 *** | 0.15 (0.03), z = 5.65, p < 0.001 *** | | | | | | | | | | | | 0.05 (0.03), z = 1.83, p = 0.067 | | | | | | | | 0.696 | 0.32, p < 0.001 *** | 103.56 | 9.69 |
| 33 | habitat_area + str_connectivity_2800m | 3.19 (0.03), z = 119.21, p < 0.001 *** | 0.15 (0.03), z = 5.52, p < 0.001 *** | | | | | | | | | | | | | 0.04 (0.03), z = 1.61, p = 0.107 | | | | | | | 0.685 | 0.33, p < 0.001 *** | 104.23 | 10.36 |
| 34 | habitat_area + str_connectivity_3000m | 3.19 (0.03), z = 119.68, p < 0.001 *** | 0.15 (0.03), z = 5.55, p < 0.001 *** | | | | | | | | | | | | | | 0.05 (0.03), z = 1.67, p = 0.095 | | | | | | 0.688 | 0.33, p < 0.001 *** | 104.07 | 10.2 |
| 35 | habitat_area + str_connectivity_3200m | 3.19 (0.03), z = 120.54, p < 0.001 *** | 0.15 (0.03), z = 5.56, p < 0.001 *** | | | | | | | | | | | | | | | 0.05 (0.03), z = 1.76, p = 0.078 | | | | | 0.692 | 0.32, p < 0.001 *** | 103.79 | 9.92 |
| 36 | habitat_area + str_connectivity_3400m | 3.19 (0.03), z = 120.75, p < 0.001 *** | 0.15 (0.03), z = 5.51, p < 0.001 *** | | | | | | | | | | | | | | | | 0.05 (0.03), z = 1.79, p = 0.074 | | | | 0.693 | 0.32, p < 0.001 *** | 103.72 | 9.85 |
| 37 | habitat_area + str_connectivity_3600m | 3.19 (0.03), z = 120.72, p < 0.001 *** | 0.15 (0.03), z = 5.40, p < 0.001 *** | | | | | | | | | | | | | | | | | 0.05 (0.03), z = 1.78, p = 0.075 | | | 0.693 | 0.32, p < 0.001 *** | 103.73 | 9.86 |
| 38 | habitat_area + str_connectivity_3800m | 3.19 (0.03), z = 120.77, p < 0.001 *** | 0.15 (0.03), z = 5.30, p < 0.001 *** | | | | | | | | | | | | | | | | | | 0.05 (0.03), z = 1.79, p = 0.074 | | 0.693 | 0.32, p < 0.001 *** | 103.71 | 9.84 |
| 39 | habitat_area + str_connectivity_4000m | 3.19 (0.03), z = 121.05, p < 0.001 *** | 0.15 (0.03), z = 5.32, p < 0.001 *** | | | | | | | | | | | | | | | | | | | 0.05 (0.03), z = 1.82, p = 0.069 | 0.695 | 0.32, p < 0.001 *** | 103.62 | 9.75 |
| 40 | shannon_index + str_connectivity_600m | 3.21 (0.05), z = 69.95, p < 0.001 *** | | -0.03 (0.05), z = -0.58, p = 0.560 | -0.03 (0.05), z = -0.70, p = 0.482 | | | | | | | | | | | | | | | | | | 0.039 | 0.97, p = 0.913 | 125.33 | 31.46 |
| 41 | shannon_index + str_connectivity_800m | 3.21 (0.04), z = 72.17, p < 0.001 *** | | -0.03 (0.05), z = -0.64, p = 0.519 | | -0.06 (0.05), z = -1.35, p = 0.176 | | | | | | | | | | | | | | | | | 0.101 | 0.92, p = 0.665 | 124.06 | 30.19 |
| 42 | shannon_index + str_connectivity_1000m | 3.21 (0.05), z = 69.72, p < 0.001 *** | | -0.02 (0.05), z = -0.52, p = 0.600 | | | -0.03 (0.05), z = -0.60, p = 0.549 | | | | | | | | | | | | | | | | 0.032 | 0.98, p = 0.927 | 125.47 | 31.6 |
| 43 | shannon_index + str_connectivity_1200m | 3.21 (0.05), z = 70.86, p < 0.001 *** | | -0.03 (0.05), z = -0.63, p = 0.528 | | | | 0.05 (0.05), z = 1.02, p = 0.306 | | | | | | | | | | | | | | | 0.065 | 0.95, p = 0.855 | 124.81 | 30.94 |
| 44 | shannon_index + str_connectivity_1400m | 3.20 (0.04), z = 76.90, p < 0.001 *** | | -0.04 (0.04), z = -1.03, p = 0.304 | | | | | 0.10 (0.04), z = 2.21, p = 0.027 * | | | | | | | | | | | | | | 0.214 | 0.8, p = 0.386 | 121.51 | 27.64 |
| 45 | shannon_index + str_connectivity_1600m | 3.20 (0.04), z = 89.15, p < 0.001 *** | | -0.08 (0.04), z = -1.97, p = 0.049 * | | | | | | 0.14 (0.04), z = 3.66, p < 0.001 *** | | | | | | | | | | | | | 0.425 | 0.6, p = 0.006 ** | 115.61 | 21.74 |
| 46 | shannon_index + str_connectivity_1800m | 3.20 (0.03), z = 95.41, p < 0.001 *** | | -0.10 (0.04), z = -2.63, p = 0.009 ** | | | | | | | 0.17 (0.04), z = 4.25, p < 0.001 *** | | | | | | | | | | | | 0.501 | 0.52, p < 0.001 *** | 112.92 | 19.05 |
| 47 | shannon_index + str_connectivity_2000m | 3.20 (0.04), z = 84.12, p < 0.001 *** | | -0.09 (0.04), z = -2.08, p = 0.038 * | | | | | | | | 0.14 (0.04), z = 3.12, p = 0.002 ** | | | | | | | | | | | 0.35 | 0.67, p = 0.012 * | 117.92 | 24.05 |
| 48 | shannon_index + str_connectivity_2200m | 3.20 (0.04), z = 79.78, p < 0.001 *** | | -0.06 (0.04), z = -1.44, p = 0.151 | | | | | | | | | 0.11 (0.04), z = 2.59, p = 0.009 ** | | | | | | | | | | 0.273 | 0.75, p = 0.164 | 120.04 | 26.17 |
| 49 | shannon_index + str_connectivity_2400m | 3.20 (0.04), z = 77.56, p < 0.001 *** | | -0.03 (0.04), z = -0.73, p = 0.465 | | | | | | | | | | 0.10 (0.04), z = 2.30, p = 0.021 * | | | | | | | | | 0.228 | 0.8, p = 0.304 | 121.17 | 27.3 |
| 50 | shannon_index + str_connectivity_2600m | 3.20 (0.04), z = 76.59, p < 0.001 *** | | -0.02 (0.04), z = -0.53, p = 0.593 | | | | | | | | | | | 0.09 (0.04), z = 2.17, p = 0.030 * | | | | | | | | 0.207 | 0.81, p = 0.321 | 121.68 | 27.81 |
| 51 | shannon_index + str_connectivity_2800m | 3.20 (0.04), z = 76.25, p < 0.001 *** | | -0.02 (0.04), z = -0.47, p = 0.639 | | | | | | | | | | | | 0.09 (0.04), z = 2.12, p = 0.034 * | | | | | | | 0.2 | 0.82, p = 0.324 | 121.85 | 27.98 |
| 52 | shannon_index + str_connectivity_3000m | 3.20 (0.04), z = 76.25, p < 0.001 *** | | -0.02 (0.04), z = -0.45, p = 0.655 | | | | | | | | | | | | | 0.09 (0.04), z = 2.12, p = 0.034 * | | | | | | 0.2 | 0.82, p = 0.329 | 121.85 | 27.98 |
| 53 | shannon_index + str_connectivity_3200m | 3.20 (0.04), z = 76.72, p < 0.001 *** | | -0.02 (0.04), z = -0.46, p = 0.646 | | | | | | | | | | | | | | 0.09 (0.04), z = 2.19, p = 0.029 * | | | | | 0.21 | 0.81, p = 0.296 | 121.61 | 27.74 |
| 54 | shannon_index + str_connectivity_3400m | 3.20 (0.04), z = 77.39, p < 0.001 *** | | -0.02 (0.04), z = -0.54, p = 0.588 | | | | | | | | | | | | | | | 0.09 (0.04), z = 2.29, p = 0.022 * | | | | 0.225 | 0.79, p = 0.244 | 121.25 | 27.38 |
| 55 | shannon_index + str_connectivity_3600m | 3.20 (0.04), z = 78.41, p < 0.001 *** | | -0.03 (0.04), z = -0.62, p = 0.535 | | | | | | | | | | | | | | | | 0.10 (0.04), z = 2.44, p = 0.015 * | | | 0.246 | 0.77, p = 0.19 | 120.73 | 26.86 |
| 56 | shannon_index + str_connectivity_3800m | 3.20 (0.04), z = 79.32, p < 0.001 *** | | -0.03 (0.04), z = -0.63, p = 0.531 | | | | | | | | | | | | | | | | | 0.10 (0.04), z = 2.56, p = 0.010 * | | 0.264 | 0.75, p = 0.155 | 120.27 | 26.4 |
| 57 | shannon_index + str_connectivity_4000m | 3.20 (0.04), z = 79.57, p < 0.001 *** | | -0.03 (0.04), z = -0.71, p = 0.476 | | | | | | | | | | | | | | | | | | 0.10 (0.04), z = 2.59, p = 0.010 ** | 0.269 | 0.75, p = 0.153 | 120.14 | 26.27 |
| 58 | habitat_area + shannon_index + str_connectivity_600m | 3.19 (0.03), z = 121.65, p < 0.001 *** | 0.18 (0.03), z = 6.41, p < 0.001 *** | 0.03 (0.03), z = 0.92, p = 0.359 | -0.04 (0.03), z = -1.54, p = 0.125 | | | | | | | | | | | | | | | | | | 0.698 | 0.32, p < 0.001 *** | 107.2 | 13.33 |
| 59 | habitat_area + shannon_index + str_connectivity_800m | 3.19 (0.03), z = 118.67, p < 0.001 *** | 0.17 (0.03), z = 5.91, p < 0.001 *** | 0.03 (0.03), z = 0.89, p = 0.372 | | -0.03 (0.03), z = -1.15, p = 0.249 | | | | | | | | | | | | | | | | | 0.682 | 0.33, p < 0.001 *** | 108.16 | 14.29 |
| 60 | habitat_area + shannon_index + str_connectivity_1000m | 3.19 (0.03), z = 115.08, p < 0.001 *** | 0.17 (0.03), z = 5.94, p < 0.001 *** | 0.03 (0.03), z = 0.99, p = 0.324 | | | -0.01 (0.03), z = -0.27, p = 0.784 | | | | | | | | | | | | | | | | 0.661 | 0.35, p < 0.001 *** | 109.37 | 15.5 |
| 61 | habitat_area + shannon_index + str_connectivity_1200m | 3.19 (0.03), z = 116.85, p < 0.001 *** | 0.17 (0.03), z = 5.90, p < 0.001 *** | 0.03 (0.03), z = 0.89, p = 0.376 | | | | 0.02 (0.03), z = 0.82, p = 0.410 | | | | | | | | | | | | | | | 0.671 | 0.35, p < 0.001 *** | 108.78 | 14.91 |
| 62 | habitat_area + shannon_index + str_connectivity_1400m | 3.19 (0.03), z = 127.62, p < 0.001 *** | 0.16 (0.03), z = 5.93, p < 0.001 *** | 0.01 (0.03), z = 0.54, p = 0.590 | | | | | 0.06 (0.03), z = 2.14, p = 0.032 * | | | | | | | | | | | | | | 0.726 | 0.29, p < 0.001 *** | 105.35 | 11.48 |
| 63 | habitat_area + shannon_index + str_connectivity_1600m | 3.19 (0.02), z = 151.26, p < 0.001 *** | 0.14 (0.02), z = 6.14, p < 0.001 *** | -0.01 (0.02), z = -0.36, p = 0.718 | | | | | | 0.09 (0.02), z = 3.79, p < 0.001 *** | | | | | | | | | | | | | 0.807 | 0.21, p < 0.001 *** | 98.76 | 4.89 |
| 64 | habitat_area + shannon_index + str_connectivity_1800m | 3.19 (0.02), z = 160.28, p < 0.001 *** | 0.14 (0.02), z = 6.05, p < 0.001 *** | -0.03 (0.03), z = -1.07, p = 0.284 | | | | | | | 0.11 (0.03), z = 4.30, p < 0.001 *** | | | | | | | | | | | | 0.829 | 0.18, p < 0.001 *** | 96.51 | 2.64 |
| 65 | habitat_area + shannon_index + str_connectivity_2000m | 3.19 (0.02), z = 148.46, p < 0.001 *** | 0.15 (0.02), z = 6.54, p < 0.001 *** | -0.02 (0.03), z = -0.74, p = 0.457 | | | | | | | | 0.10 (0.03), z = 3.63, p < 0.001 *** | | | | | | | | | | | 0.8 | 0.21, p < 0.001 *** | 99.46 | 5.59 |
| 66 | habitat_area + shannon_index + str_connectivity_2200m | 3.19 (0.02), z = 134.89, p < 0.001 *** | 0.16 (0.03), z = 6.17, p < 0.001 *** | 0.00 (0.03), z = 0.08, p = 0.937 | | | | | | | | | 0.07 (0.03), z = 2.72, p = 0.007 ** | | | | | | | | | | 0.756 | 0.26, p < 0.001 *** | 103.18 | 9.31 |
| 67 | habitat_area + shannon_index + str_connectivity_2400m | 3.19 (0.02), z = 128.06, p < 0.001 *** | 0.16 (0.03), z = 5.97, p < 0.001 *** | 0.02 (0.03), z = 0.83, p = 0.407 | | | | | | | | | | 0.06 (0.03), z = 2.18, p = 0.029 * | | | | | | | | | 0.728 | 0.28, p < 0.001 *** | 105.19 | 11.32 |
| 68 | habitat_area + shannon_index + str_connectivity_2600m | 3.19 (0.03), z = 124.12, p < 0.001 *** | 0.16 (0.03), z = 5.80, p < 0.001 *** | 0.03 (0.03), z = 0.97, p = 0.333 | | | | | | | | | | | 0.05 (0.03), z = 1.82, p = 0.069 | | | | | | | | 0.71 | 0.3, p < 0.001 *** | 106.41 | 12.54 |
| 69 | habitat_area + shannon_index + str_connectivity_2800m | 3.19 (0.03), z = 122.16, p < 0.001 *** | 0.16 (0.03), z = 5.69, p < 0.001 *** | 0.03 (0.03), z = 0.99, p = 0.322 | | | | | | | | | | | | 0.04 (0.03), z = 1.61, p = 0.108 | | | | | | | 0.701 | 0.31, p < 0.001 *** | 107.03 | 13.16 |
| 70 | habitat_area + shannon_index + str_connectivity_3000m | 3.19 (0.03), z = 122.77, p < 0.001 *** | 0.16 (0.03), z = 5.73, p < 0.001 *** | 0.03 (0.03), z = 1.01, p = 0.312 | | | | | | | | | | | | | 0.04 (0.03), z = 1.68, p = 0.093 | | | | | | 0.704 | 0.31, p < 0.001 *** | 106.83 | 12.96 |
| 71 | habitat_area + shannon_index + str_connectivity_3200m | 3.19 (0.03), z = 123.62, p < 0.001 *** | 0.16 (0.03), z = 5.73, p < 0.001 *** | 0.03 (0.03), z = 1.01, p = 0.314 | | | | | | | | | | | | | | 0.05 (0.03), z = 1.77, p = 0.077 | | | | | 0.708 | 0.31, p < 0.001 *** | 106.56 | 12.69 |
| 72 | habitat_area + shannon_index + str_connectivity_3400m | 3.19 (0.03), z = 123.40, p < 0.001 *** | 0.16 (0.03), z = 5.63, p < 0.001 *** | 0.03 (0.03), z = 0.93, p = 0.352 | | | | | | | | | | | | | | | 0.05 (0.03), z = 1.75, p = 0.081 | | | | 0.707 | 0.31, p < 0.001 *** | 106.64 | 12.77 |
| 73 | habitat_area + shannon_index + str_connectivity_3600m | 3.19 (0.03), z = 122.95, p < 0.001 *** | 0.16 (0.03), z = 5.48, p < 0.001 *** | 0.02 (0.03), z = 0.85, p = 0.394 | | | | | | | | | | | | | | | | 0.05 (0.03), z = 1.70, p = 0.089 | | | 0.705 | 0.31, p < 0.001 *** | 106.78 | 12.91 |
| 74 | habitat_area + shannon_index + str_connectivity_3800m | 3.19 (0.03), z = 122.93, p < 0.001 *** | 0.16 (0.03), z = 5.37, p < 0.001 *** | 0.02 (0.03), z = 0.84, p = 0.401 | | | | | | | | | | | | | | | | | 0.05 (0.03), z = 1.70, p = 0.090 | | 0.705 | 0.31, p < 0.001 *** | 106.78 | 12.91 |
| 75 | habitat_area + shannon_index + str_connectivity_4000m | 3.19 (0.03), z = 122.88, p < 0.001 *** | 0.16 (0.03), z = 5.34, p < 0.001 *** | 0.02 (0.03), z = 0.77, p = 0.441 | | | | | | | | | | | | | | | | | | 0.05 (0.03), z = 1.69, p = 0.091 | 0.704 | 0.31, p < 0.001 *** | 106.8 | 12.93 |
