## Supplementary material for "How to build dense cities without jeopardizing biodiversity? Insights from bird observations in Gothenburg, Sweden": Table S3

Table S3. Results of the fitted PERMANOVA candidate models examining the relationship between species composition and three environmental variables (habitat area, Shannon index, and structural connectivity at multiple spatial scales). Each row represents a candidate model including one, two, or three environmental variables. For each variable included in a model, the marginal $R^2$ and permutation-based p-value (with significance level) are reported. The table also reports the residual $R^2$ for every model. Variables not included in a model are left blank. Significance levels are indicated as: $p < 0.05$ = *, $p < 0.01$ = **, $p < 0.001$ = ***.

| Model ID | Model | habitat_area | shannon_index | str_connectivity_600m | str_connectivity_800m | str_connectivity_1000m | str_connectivity_1200m | str_connectivity_1400m | str_connectivity_1600m | str_connectivity_1800m | str_connectivity_2000m | str_connectivity_2200m | str_connectivity_2400m | str_connectivity_2600m | str_connectivity_2800m | str_connectivity_3000m | str_connectivity_3200m | str_connectivity_3400m | str_connectivity_3600m | str_connectivity_3800m | str_connectivity_4000m | Residual R2 |
|---|---|---|---|---|---|---|---|---|---|---|---|---|---|---|---|---|---|---|---|---|---|---|
| 1 | habitat_area | R2 = 0.217, p < 0.001 *** | | | | | | | | | | | | | | | | | | | | 0.783 |
| 2 | shannon_index | | R2 = 0.029, p = 0.888 | | | | | | | | | | | | | | | | | | | 0.971 |
| 3 | str_connectivity_600m | | | R2 = 0.037, p = 0.773 | | | | | | | | | | | | | | | | | | 0.963 |
| 4 | str_connectivity_800m | | | | R2 = 0.029, p = 0.891 | | | | | | | | | | | | | | | | | 0.971 |
| 5 | str_connectivity_1000m | | | | | R2 = 0.034, p = 0.808 | | | | | | | | | | | | | | | | 0.966 |
| 6 | str_connectivity_1200m | | | | | | R2 = 0.067, p = 0.259 | | | | | | | | | | | | | | | 0.933 |
| 7 | str_connectivity_1400m | | | | | | | R2 = 0.096, p = 0.072 | | | | | | | | | | | | | | 0.904 |
| 8 | str_connectivity_1600m | | | | | | | | R2 = 0.102, p = 0.058 | | | | | | | | | | | | | 0.898 |
| 9 | str_connectivity_1800m | | | | | | | | | R2 = 0.101, p = 0.06 | | | | | | | | | | | | 0.899 |
| 10 | str_connectivity_2000m | | | | | | | | | | R2 = 0.091, p = 0.091 | | | | | | | | | | | 0.909 |
| 11 | str_connectivity_2200m | | | | | | | | | | | R2 = 0.101, p = 0.063 | | | | | | | | | | 0.899 |
| 12 | str_connectivity_2400m | | | | | | | | | | | | R2 = 0.112, p = 0.039 * | | | | | | | | | 0.888 |
| 13 | str_connectivity_2600m | | | | | | | | | | | | | R2 = 0.112, p = 0.039 * | | | | | | | | 0.888 |
| 14 | str_connectivity_2800m | | | | | | | | | | | | | | R2 = 0.113, p = 0.038 * | | | | | | | 0.887 |
| 15 | str_connectivity_3000m | | | | | | | | | | | | | | | R2 = 0.118, p = 0.031 * | | | | | | 0.882 |
| 16 | str_connectivity_3200m | | | | | | | | | | | | | | | | R2 = 0.124, p = 0.024 * | | | | | 0.876 |
| 17 | str_connectivity_3400m | | | | | | | | | | | | | | | | | R2 = 0.128, p = 0.02 * | | | | 0.872 |
| 18 | str_connectivity_3600m | | | | | | | | | | | | | | | | | | R2 = 0.134, p = 0.016 * | | | 0.866 |
| 19 | str_connectivity_3800m | | | | | | | | | | | | | | | | | | | R2 = 0.140, p = 0.012 * | | 0.86 |
| 20 | str_connectivity_4000m | | | | | | | | | | | | | | | | | | | | R2 = 0.142, p = 0.011 * | 0.858 |
| 21 | habitat_area + shannon_index | R2 = 0.204, p < 0.001 *** | R2 = 0.016, p = 0.976 | | | | | | | | | | | | | | | | | | | 0.766 |
| 22 | habitat_area + str_connectivity_600m | R2 = 0.221, p < 0.001 *** | | R2 = 0.041, p = 0.531 | | | | | | | | | | | | | | | | | | 0.742 |
| 23 | habitat_area + str_connectivity_800m | R2 = 0.211, p < 0.001 *** | | | R2 = 0.023, p = 0.913 | | | | | | | | | | | | | | | | | 0.76 |
| 24 | habitat_area + str_connectivity_1000m | R2 = 0.224, p < 0.001 *** | | | | R2 = 0.041, p = 0.52 | | | | | | | | | | | | | | | | 0.742 |
| 25 | habitat_area + str_connectivity_1200m | R2 = 0.207, p < 0.001 *** | | | | | R2 = 0.056, p = 0.252 | | | | | | | | | | | | | | | 0.727 |
| 26 | habitat_area + str_connectivity_1400m | R2 = 0.193, p < 0.001 *** | | | | | | R2 = 0.071, p = 0.11 | | | | | | | | | | | | | | 0.711 |
| 27 | habitat_area + str_connectivity_1600m | R2 = 0.182, p < 0.001 *** | | | | | | | R2 = 0.066, p = 0.147 | | | | | | | | | | | | | 0.716 |
| 28 | habitat_area + str_connectivity_1800m | R2 = 0.179, p < 0.001 *** | | | | | | | | R2 = 0.063, p = 0.177 | | | | | | | | | | | | 0.72 |
| 29 | habitat_area + str_connectivity_2000m | R2 = 0.198, p < 0.001 *** | | | | | | | | | R2 = 0.072, p = 0.111 | | | | | | | | | | | 0.711 |
| 30 | habitat_area + str_connectivity_2200m | R2 = 0.189, p < 0.001 *** | | | | | | | | | | R2 = 0.072, p = 0.11 | | | | | | | | | | 0.711 |
| 31 | habitat_area + str_connectivity_2400m | R2 = 0.172, p = 0.001 ** | | | | | | | | | | | R2 = 0.066, p = 0.146 | | | | | | | | | 0.716 |
| 32 | habitat_area + str_connectivity_2600m | R2 = 0.168, p = 0.001 ** | | | | | | | | | | | | R2 = 0.063, p = 0.173 | | | | | | | | 0.72 |
| 33 | habitat_area + str_connectivity_2800m | R2 = 0.165, p = 0.002 ** | | | | | | | | | | | | | R2 = 0.060, p = 0.197 | | | | | | | 0.722 |
| 34 | habitat_area + str_connectivity_3000m | R2 = 0.163, p = 0.002 ** | | | | | | | | | | | | | | R2 = 0.063, p = 0.168 | | | | | | 0.719 |
| 35 | habitat_area + str_connectivity_3200m | R2 = 0.161, p = 0.002 ** | | | | | | | | | | | | | | | R2 = 0.068, p = 0.131 | | | | | 0.714 |
| 36 | habitat_area + str_connectivity_3400m | R2 = 0.159, p = 0.002 ** | | | | | | | | | | | | | | | | R2 = 0.070, p = 0.122 | | | | 0.713 |
| 37 | habitat_area + str_connectivity_3600m | R2 = 0.153, p = 0.003 ** | | | | | | | | | | | | | | | | | R2 = 0.069, p = 0.128 | | | 0.714 |
| 38 | habitat_area + str_connectivity_3800m | R2 = 0.146, p = 0.004 ** | | | | | | | | | | | | | | | | | | R2 = 0.069, p = 0.128 | | 0.714 |
| 39 | habitat_area + str_connectivity_4000m | R2 = 0.145, p = 0.004 ** | | | | | | | | | | | | | | | | | | | R2 = 0.070, p = 0.124 | 0.713 |
| 40 | shannon_index + str_connectivity_600m | | R2 = 0.030, p = 0.89 | R2 = 0.037, p = 0.783 | | | | | | | | | | | | | | | | | | 0.933 |
| 41 | shannon_index + str_connectivity_800m | | R2 = 0.030, p = 0.887 | | R2 = 0.030, p = 0.89 | | | | | | | | | | | | | | | | | 0.941 |
| 42 | shannon_index + str_connectivity_1000m | | R2 = 0.029, p = 0.897 | | | R2 = 0.034, p = 0.825 | | | | | | | | | | | | | | | | 0.936 |
| 43 | shannon_index + str_connectivity_1200m | | R2 = 0.034, p = 0.806 | | | | R2 = 0.072, p = 0.227 | | | | | | | | | | | | | | | 0.899 |
| 44 | shannon_index + str_connectivity_1400m | | R2 = 0.044, p = 0.579 | | | | | R2 = 0.111, p = 0.044 * | | | | | | | | | | | | | | 0.86 |
| 45 | shannon_index + str_connectivity_1600m | | R2 = 0.063, p = 0.26 | | | | | | R2 = 0.136, p = 0.016 * | | | | | | | | | | | | | 0.835 |
| 46 | shannon_index + str_connectivity_1800m | | R2 = 0.085, p = 0.1 | | | | | | | R2 = 0.157, p = 0.006 ** | | | | | | | | | | | | 0.814 |
| 47 | shannon_index + str_connectivity_2000m | | R2 = 0.088, p = 0.091 | | | | | | | | R2 = 0.150, p = 0.008 ** | | | | | | | | | | | 0.82 |
| 48 | shannon_index + str_connectivity_2200m | | R2 = 0.065, p = 0.237 | | | | | | | | | R2 = 0.137, p = 0.016 * | | | | | | | | | | 0.834 |
| 49 | shannon_index + str_connectivity_2400m | | R2 = 0.036, p = 0.743 | | | | | | | | | | R2 = 0.119, p = 0.034 * | | | | | | | | | 0.852 |
| 50 | shannon_index + str_connectivity_2600m | | R2 = 0.029, p = 0.871 | | | | | | | | | | | R2 = 0.112, p = 0.045 * | | | | | | | | 0.859 |
| 51 | shannon_index + str_connectivity_2800m | | R2 = 0.026, p = 0.905 | | | | | | | | | | | | R2 = 0.110, p = 0.048 * | | | | | | | 0.861 |
| 52 | shannon_index + str_connectivity_3000m | | R2 = 0.026, p = 0.913 | | | | | | | | | | | | | R2 = 0.114, p = 0.04 * | | | | | | 0.857 |
| 53 | shannon_index + str_connectivity_3200m | | R2 = 0.026, p = 0.906 | | | | | | | | | | | | | | R2 = 0.121, p = 0.031 * | | | | | 0.85 |
| 54 | shannon_index + str_connectivity_3400m | | R2 = 0.029, p = 0.866 | | | | | | | | | | | | | | | R2 = 0.127, p = 0.024 * | | | | 0.844 |
| 55 | shannon_index + str_connectivity_3600m | | R2 = 0.031, p = 0.823 | | | | | | | | | | | | | | | | R2 = 0.135, p = 0.017 * | | | 0.835 |
| 56 | shannon_index + str_connectivity_3800m | | R2 = 0.031, p = 0.822 | | | | | | | | | | | | | | | | | R2 = 0.142, p = 0.013 * | | 0.829 |
| 57 | shannon_index + str_connectivity_4000m | | R2 = 0.033, p = 0.769 | | | | | | | | | | | | | | | | | | R2 = 0.146, p = 0.011 * | 0.825 |
| 58 | habitat_area + shannon_index + str_connectivity_600m | R2 = 0.206, p < 0.001 *** | R2 = 0.015, p = 0.985 | R2 = 0.039, p = 0.604 | | | | | | | | | | | | | | | | | | 0.727 |
| 59 | habitat_area + shannon_index + str_connectivity_800m | R2 = 0.196, p < 0.001 *** | R2 = 0.015, p = 0.984 | | R2 = 0.022, p = 0.939 | | | | | | | | | | | | | | | | | 0.745 |
| 60 | habitat_area + shannon_index + str_connectivity_1000m | R2 = 0.211, p < 0.001 *** | R2 = 0.017, p = 0.974 | | | R2 = 0.041, p = 0.546 | | | | | | | | | | | | | | | | 0.725 |
| 61 | habitat_area + shannon_index + str_connectivity_1200m | R2 = 0.189, p < 0.001 *** | R2 = 0.016, p = 0.974 | | | | R2 = 0.056, p = 0.281 | | | | | | | | | | | | | | | 0.71 |
| 62 | habitat_area + shannon_index + str_connectivity_1400m | R2 = 0.167, p = 0.002 ** | R2 = 0.019, p = 0.953 | | | | | R2 = 0.074, p = 0.114 | | | | | | | | | | | | | | 0.693 |
| 63 | habitat_area + shannon_index + str_connectivity_1600m | R2 = 0.138, p = 0.007 ** | R2 = 0.019, p = 0.948 | | | | | | R2 = 0.069, p = 0.143 | | | | | | | | | | | | | 0.697 |
| 64 | habitat_area + shannon_index + str_connectivity_1800m | R2 = 0.120, p = 0.015 * | R2 = 0.026, p = 0.852 | | | | | | | R2 = 0.072, p = 0.124 | | | | | | | | | | | | 0.694 |
| 65 | habitat_area + shannon_index + str_connectivity_2000m | R2 = 0.144, p = 0.004 ** | R2 = 0.034, p = 0.655 | | | | | | | | R2 = 0.089, p = 0.054 | | | | | | | | | | | 0.677 |
| 66 | habitat_area + shannon_index + str_connectivity_2200m | R2 = 0.150, p = 0.003 ** | R2 = 0.026, p = 0.834 | | | | | | | | | R2 = 0.082, p = 0.077 | | | | | | | | | | 0.684 |
| 67 | habitat_area + shannon_index + str_connectivity_2400m | R2 = 0.155, p = 0.003 ** | R2 = 0.019, p = 0.951 | | | | | | | | | | R2 = 0.069, p = 0.146 | | | | | | | | | 0.697 |
| 68 | habitat_area + shannon_index + str_connectivity_2600m | R2 = 0.157, p = 0.003 ** | R2 = 0.018, p = 0.961 | | | | | | | | | | | R2 = 0.065, p = 0.18 | | | | | | | | 0.702 |
| 69 | habitat_area + shannon_index + str_connectivity_2800m | R2 = 0.156, p = 0.003 ** | R2 = 0.017, p = 0.968 | | | | | | | | | | | | R2 = 0.061, p = 0.213 | | | | | | | 0.705 |
| 70 | habitat_area + shannon_index + str_connectivity_3000m | R2 = 0.154, p = 0.003 ** | R2 = 0.017, p = 0.971 | | | | | | | | | | | | | R2 = 0.064, p = 0.187 | | | | | | 0.703 |
| 71 | habitat_area + shannon_index + str_connectivity_3200m | R2 = 0.152, p = 0.003 ** | R2 = 0.017, p = 0.97 | | | | | | | | | | | | | | R2 = 0.069, p = 0.146 | | | | | 0.698 |
| 72 | habitat_area + shannon_index + str_connectivity_3400m | R2 = 0.148, p = 0.004 ** | R2 = 0.017, p = 0.965 | | | | | | | | | | | | | | | R2 = 0.071, p = 0.132 | | | | 0.695 |
| 73 | habitat_area + shannon_index + str_connectivity_3600m | R2 = 0.140, p = 0.005 ** | R2 = 0.018, p = 0.957 | | | | | | | | | | | | | | | | R2 = 0.071, p = 0.132 | | | 0.695 |
| 74 | habitat_area + shannon_index + str_connectivity_3800m | R2 = 0.133, p = 0.007 ** | R2 = 0.018, p = 0.956 | | | | | | | | | | | | | | | | | R2 = 0.071, p = 0.132 | | 0.695 |
| 75 | habitat_area + shannon_index + str_connectivity_4000m | R2 = 0.131, p = 0.008 ** | R2 = 0.019, p = 0.948 | | | | | | | | | | | | | | | | | | R2 = 0.072, p = 0.122 | 0.694 |
