## Supplementary figures and images for "How to build dense cities without jeopardizing biodiversity? Insights from bird observations in Gothenburg, Sweden"

### Figure S1

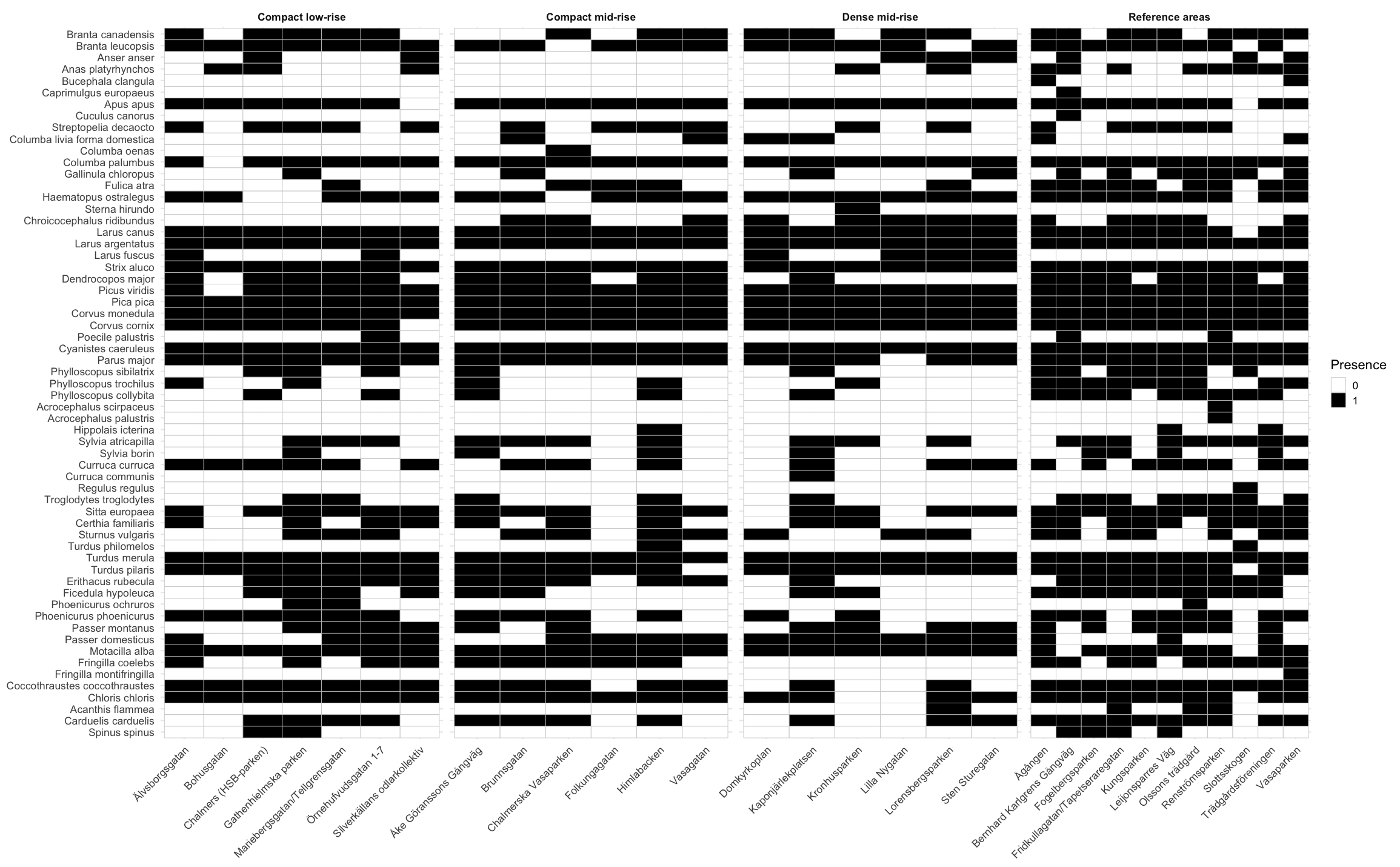

### Figure S2

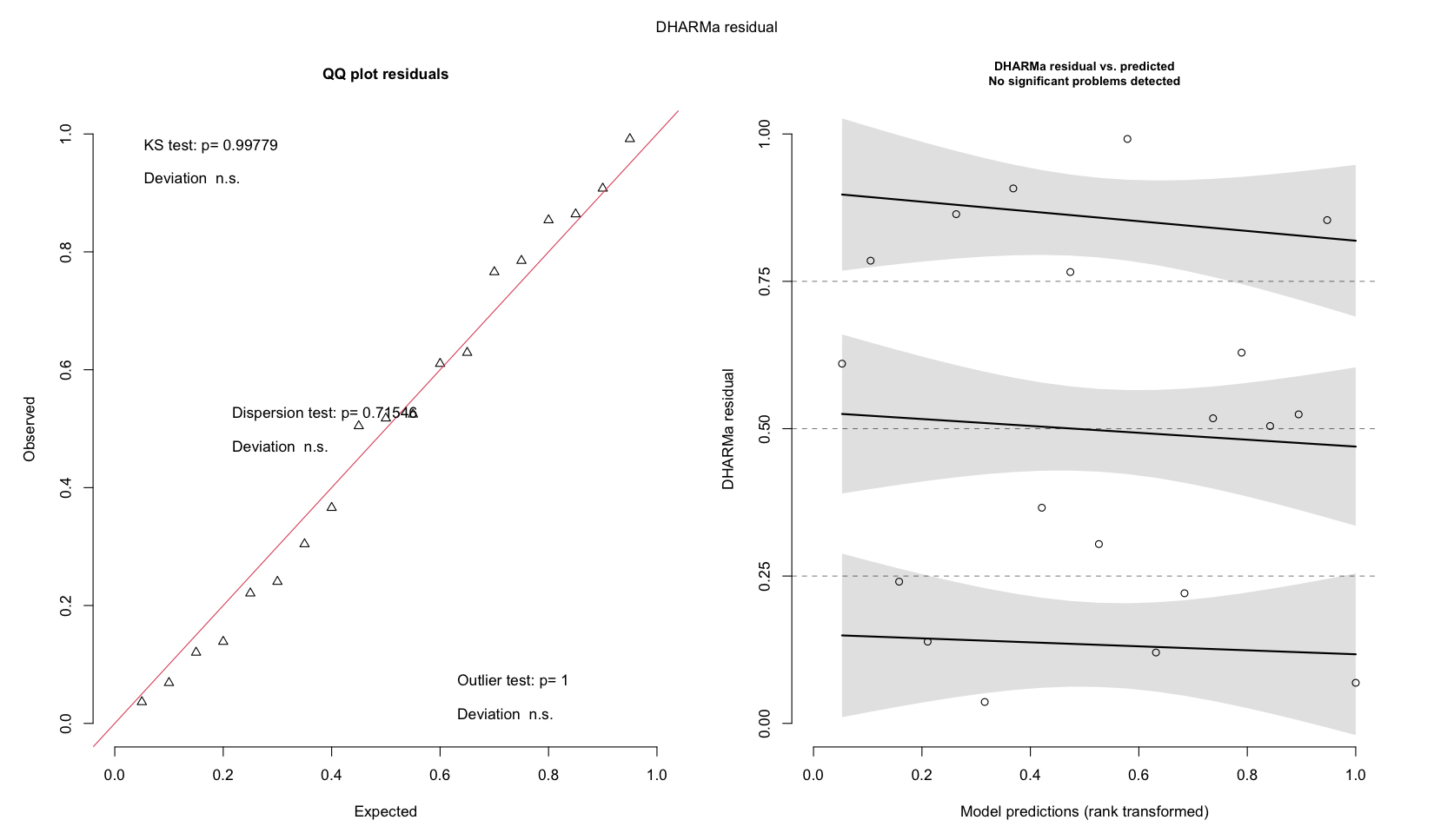

### Figure S3

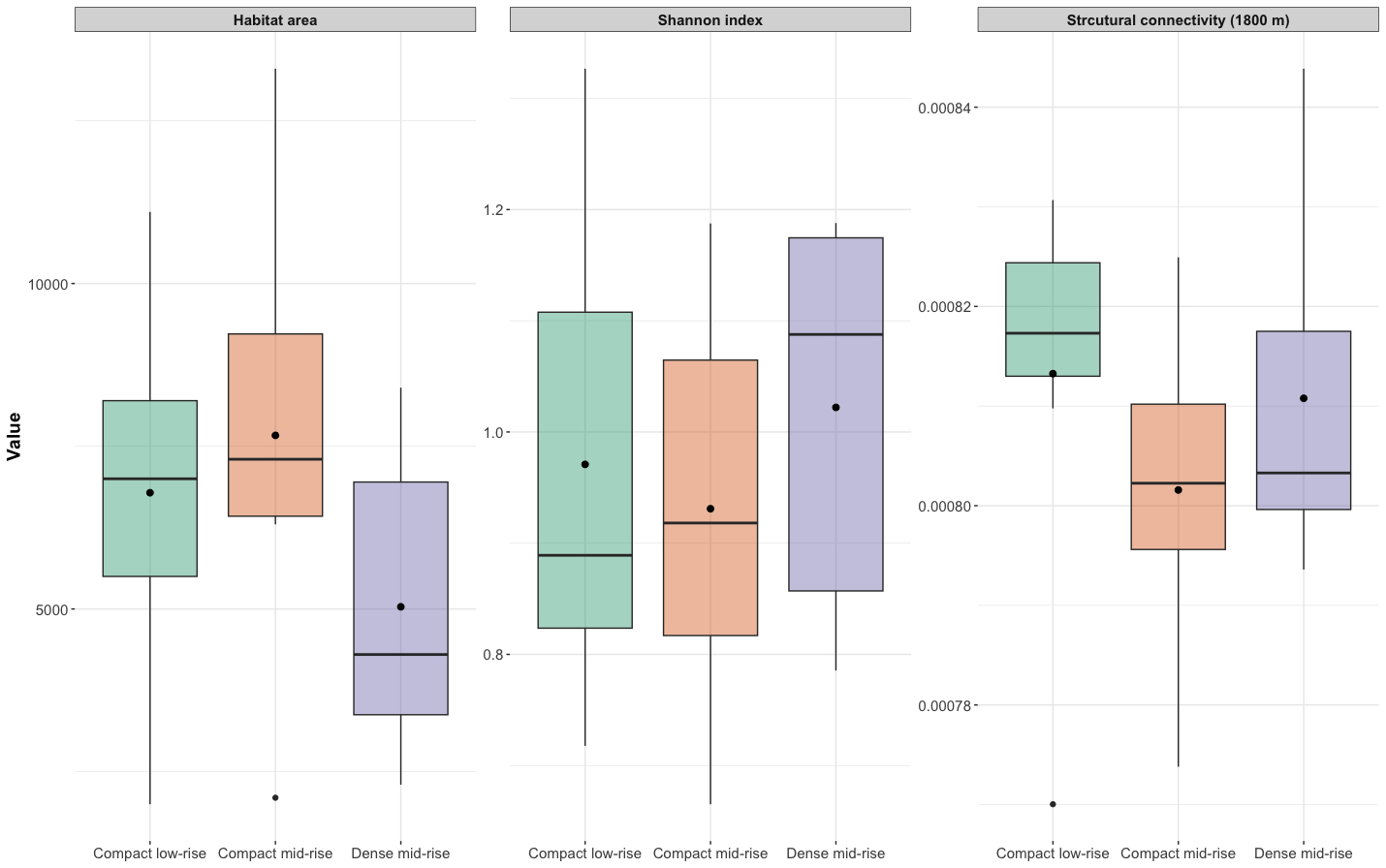

### Figure S4

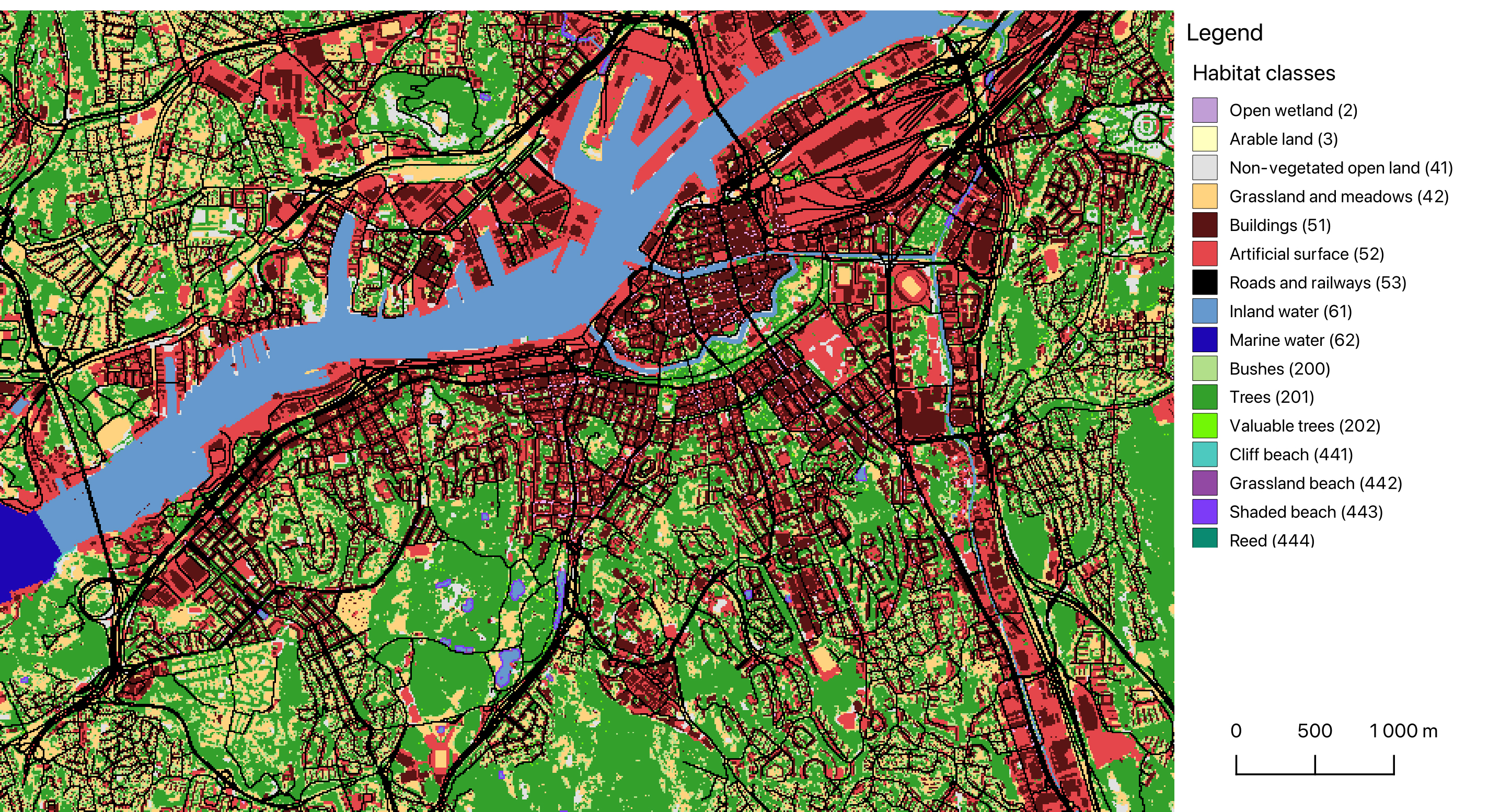

### Figure S5

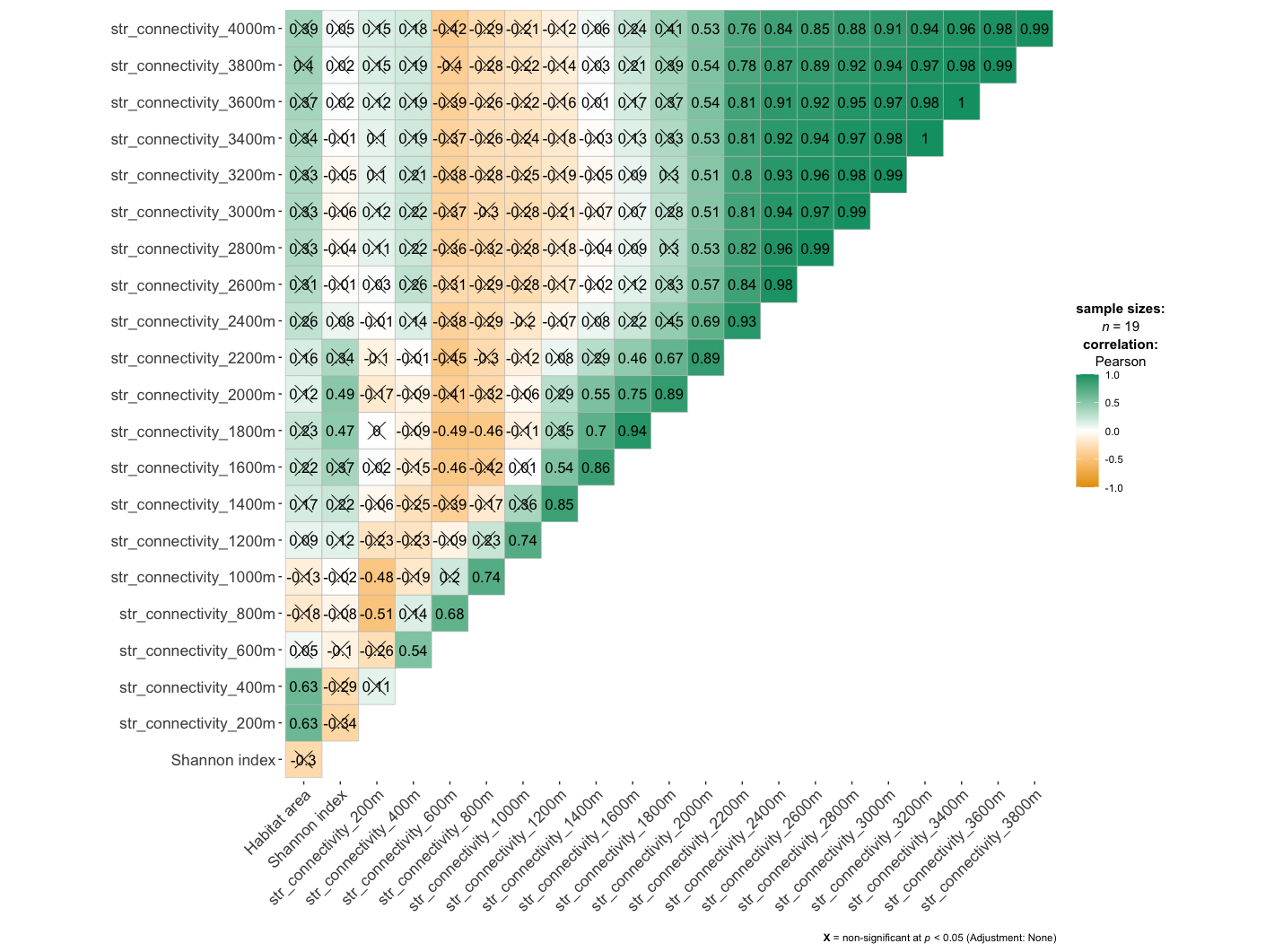
